# Genome-wide phenotypic analysis of growth, cell morphogenesis and cell cycle events in *Escherichia coli*

**DOI:** 10.1101/101832

**Authors:** Manuel Campos, Sander K. Govers, Irnov Irnov, Genevieve S. Dobihal, François Cornet, Christine Jacobs-Wagner

## Abstract

Cell size, cell growth and the cell cycle are necessarily intertwined to achieve robust bacterial replication. Yet, a comprehensive and integrated view of these fundamental processes is lacking. Here, we describe an image-based quantitative screen of the single-gene knockout collection of *Escherichia coli*, and identify many new genes involved in cell morphogenesis, population growth, nucleoid (bulk chromosome) dynamics and cell division. Functional analyses, together with high-dimensional classification, unveil new associations of morphological and cell cycle phenotypes with specific functions and pathways. Additionally, correlation analysis across ~4,000 genetic perturbations shows that growth rate is surprisingly not predictive of cell size. Growth rate was also uncorrelated with the relative timings of nucleoid separation and cell constriction. Rather, our analysis identifies scaling relationships between cell size and nucleoid size and between nucleoid size and the relative timings of nucleoid separation and cell division. These connections suggest that the nucleoid links cell morphogenesis to the cell cycle.

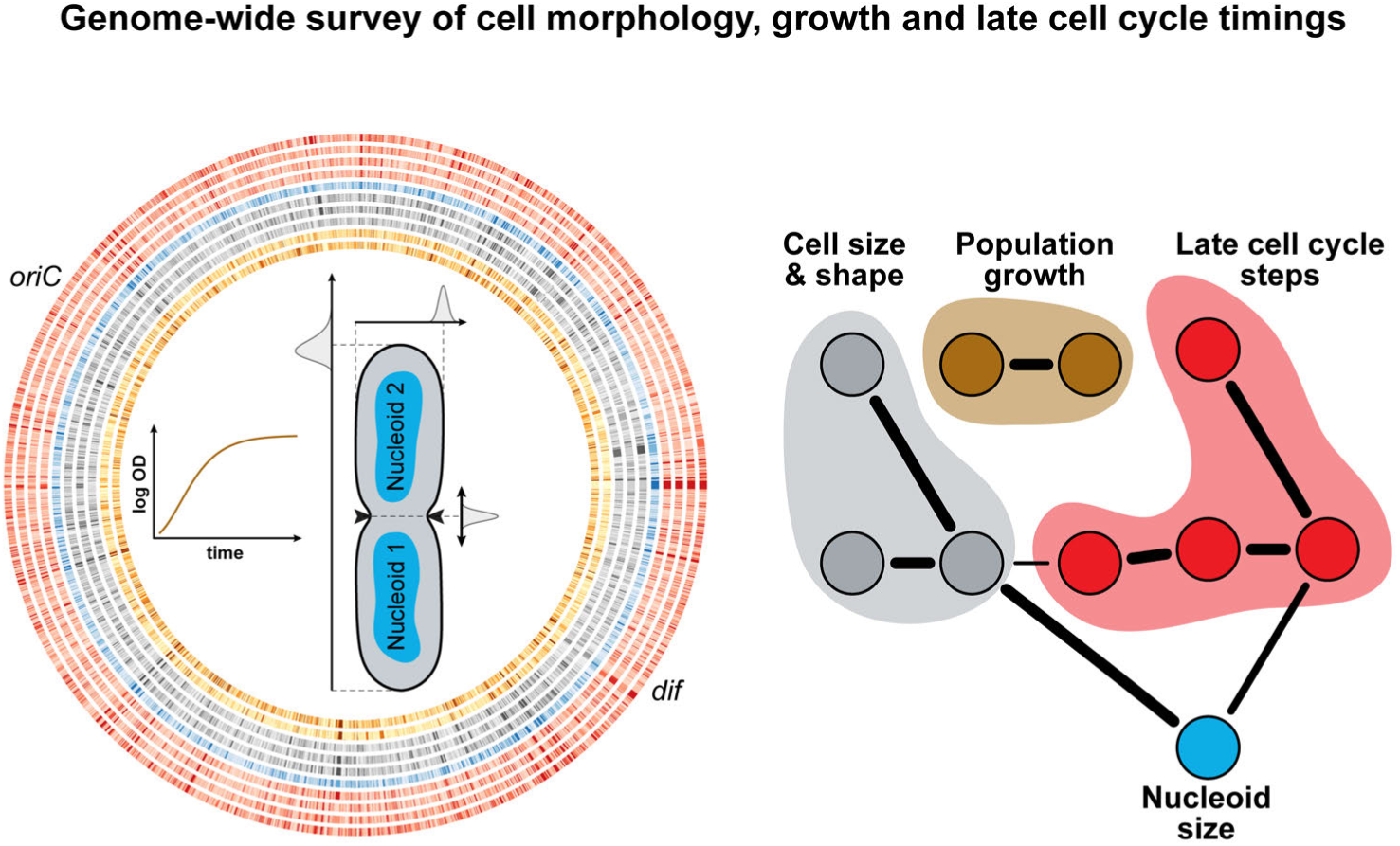

## Introduction

Cells must integrate a large variety of processes to achieve robust multiplication. Bacteria, in particular, are remarkable at proliferating, which has been key to their colonization success. During their fast-paced replication, bacterial cells must uptake and process nutrients, generate energy, build cellular components, duplicate and segregate their genetic material, couple growth and division, and maintain their shape and size, all while sensing their environment and repairing cellular damages, just to name a few important tasks. These tasks must be integrated to ensure successful cellular replication. Decades of work have garnered extensive knowledge on specific processes, genes and pathways, but we still lack a comprehensive view of the genetic determinants affecting cell morphogenesis and the cell cycle. It is also unclear how cellular activities are integrated to ensure that each division produces two viable daughter cells.

Systematic genome-wide screens, rendered possible by the creation of arrayed single-gene knock-out collections, have been successfully used to gain a more comprehensive perspective on cell morphogenesis and the cell cycle in yeast (Graml et al, 2014; Jorgensen et al, 2002; Ohya et al, 2005). Beyond the functional information gained through the mapping of phenotypes associated with the deletion of genes, genome-wide screens also provide a unique opportunity to interrogate the relationship between phenotypic features with thousands of independent genetic perturbations (Liberali et al, 2015). Here, we present a high-content, quantitative study that uses the Keio collection of *Escherichia coli* gene deletion strains (Baba et al, 2006) and combines microscopy with advanced statistical and image analysis procedures to examine the impact of each non-essential *E. coli* gene on cell morphology, cell size, growth, nucleoid (bulk chromosome) dynamics and cell constriction. In addition, we provide insight into the connectivity and empirical relationships between cell morphogenesis, growth and cell cycle events.

## Results

### High-throughput imaging and growth measurements of the *E. coli* Keio collection

To gain an understanding of the molecular relationship between growth, cell size, cell shape and specific cell cycle events, we imaged 4,227 strains of the Keio collection. This set of single gene deletion strains represents 98% of the non-essential genome (87% of the complete genome) of *E. coli* K12. The strains were grown in 96-well plates in M9 medium supplemented with 0.1% casamino acids and 0.2% glucose at 30°C. The preferred carbon source (glucose) and the casamino acids provide growth conditions that give rise to overlapping DNA replication cycles (Appendix Fig S1A). Live cells were stained with the DNA dye DAPI, and spotted on large custom-made agarose pads (48 strains per pad) prior to imaging by phase contrast and epifluorescence microscopy (Fig 1A). On average, about 360 (± 165) cells were imaged for each strain. To provide a reference, 240 replicates of the parental strain (BW25113, here referred to as WT) were also grown and imaged under the same conditions as the mutants. In parallel, using a microplate reader, we recorded the growth curves of all the strains (Fig 1A) and estimated two population-growth features. We fitted the Gompertz function to estimate the maximal growth rate (*α*_max_) and used the last hour of growth to calculate the saturating density (OD_max_) of each culture (Appendix Fig S1B). The goodness of the fits is illustrated at the time of maximal growth where the OD_600nm_ from the growth curve is highly correlated with the OD_600nm_ predicted by the fit (Appendix Fig S1C). The vast majority of strains were imaged in exponential phase at an OD_600nm_ (OD_imaging_) 4-5 times smaller than their OD_max_ (Appendix Fig S1D).

**Figure 1.**
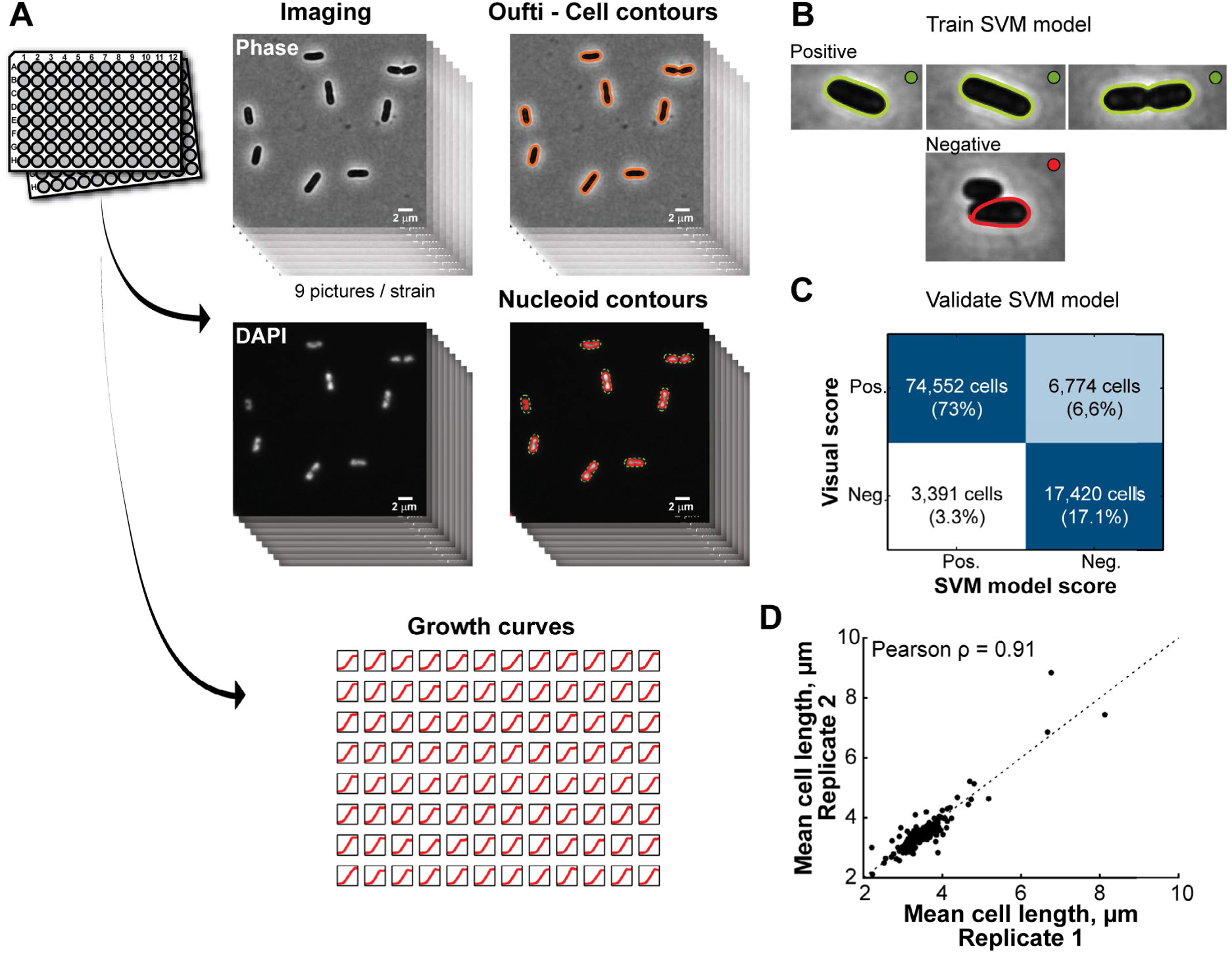
Experimental approach and reproducibility. **A.** Experimental workflow. Single gene knockout strains from the Keio collection were grown in M9 supplemented medium at 30^*◦*^C in 96-well plates. DNA was stained with DAPI prior to imaging and nine images were taken in both phase-contrast and DAPI channels. The images were then processed with MicrobeTracker and Oufti to identify the cell and nucleoid contours. In parallel, we recorded the growth curve of each imaged strain in order to extract growth parameters. **B.** A SVM model was trained via visual scoring of 43,774 cells. **C.** Confusion matrix of the SVM model based on a large validation dataset (102,137 cells), illustrating the distribution of the SVM classifier output in comparison to the visual classification. **D.** Comparison of the average cell length of 178 strains obtained from two independent 96-well cultures of the 176 most phenotypically remarkable Keio strains and 2 WT replicates.

### High-throughput dataset curation using a support vector machine

Cells and their contours were detected in an automated fashion (Sliusarenko et al, 2011). The large size (>1,500,000 detected cells) of the dataset precluded the validation of each cell contour by visual inspection. Therefore, we implemented an automated classification method based on a support vector machine (SVM) (Fan et al, 2005) to identify and discard incorrectly detected cells (Fig 1B). To generate a training dataset for the SVM model, we visually scored (positive or negative) 43,774 cell contours from the parental strain and the 419 mutants displaying the greatest deviations in cellular dimensions before data curation. The inclusion of the most aberrant mutants in the training dataset allowed us to build a versatile model that performed well on the wide range of cell sizes and shapes present in our dataset. The quality of the fit of the SVM model to the training dataset was evaluated by a 10-fold cross-validation (Hastie et al, 2009), which gave a misclassification error rate of 10%. The model was further validated on an independent dataset of 102,137 visually scored cell contours taken from the same group of WT and mutant strains. We found that our SVM model performed very well on this validation set, as shown by the high AUROC (area under the ‘receiver operating characteristic’ curve) value of 0.94 (Appendix Fig S1E). By comparing the model classification with visual scoring (Fig 1C), we found that only about 3% of cell contours in the validation set were incorrectly identified as positive by the SVM model. Importantly, these false positive cells introduced no biases in the measurement of the SVM predictor values (Appendix Fig S1F), even when considering the 419 most aberrant strains (Appendix Fig S1G). This validated SVM model was used to curate the entire dataset, retaining about 1,300,000 identified cells (291 ± 116 cells/strain). In addition, we verified the reproducibility of our experimental approach by separately imaging two independent replicates of 178 strains that included 2 copies of the parental (WT) strain and 176 mutants with severe morphological defects. We observed a Pearson correlation (*ρ*) of 0.91 for cell length (Fig 1D), indicating high reproducibility.

### Quantification of cell morphological features across the genome

With this high-quality dataset, we were able to obtain a wealth of quantitative information using the software packages MicrobeTracker and Oufti (Paintdakhi et al, 2016; Sliusarenko et al, 2011). From phase-contrast images, we measured cellular dimensions, such as length, width, perimeter, cross-sectional area, aspect ratio (width/length) and circularity (4*π* area/(perimeter)^2^). We also measured and the variability of these features by calculating their coefficient of variation (CV, the standard deviation divided by the mean). From both series of measurements, we extracted the mean and CV of additional morphological parameters, such as surface area, volume and surface-to-volume ratio. For constricted cells, we determined the relative position of division along the cell length (division ratio). Note that since the identity of the cell poles (old versus new) was unknown, randomization of cell pole identity would automatically produce a *mean* division ratio of 0.5, even for an off-center division. Therefore, measurements of mean division ratio were meaningless and not included in our analysis. However, the CV of the division ratio was included since a high CV indicated either an asymmetric division or an imprecise division site selection. In total, each strain was characterized by 19 morphological features (see Dataset EV1 for raw data). The name and abbreviation for all the features can be found in Table S1.

After taking into consideration experimental variability (see Materials and methods, Appendix Fig S2-S4,), we calculated a normalized score (*s*) for each feature and each strain (see Materials and methods). The corrected and normalized data (scores) can be found in Dataset EV2. Even with a conservative threshold of 3 standard deviations (*s* ≤-3 or ≥ 3, or absolute score |s| ≥ 3) away from the WT, a large number (874) of single gene-deletion strains were associated with one or more morphological defects (Fig 2 and Dataset EV2). This result indicates that a large fraction of the non-essential genome (i.e._~_ 20% of the unique deletion strains present in the Keio collection) directly or indirectly affect cell size and shape. Similar genomic commitment to cell size and shape was observed in budding yeast (Jorgensen et al, 2002).

**Figure 2.**
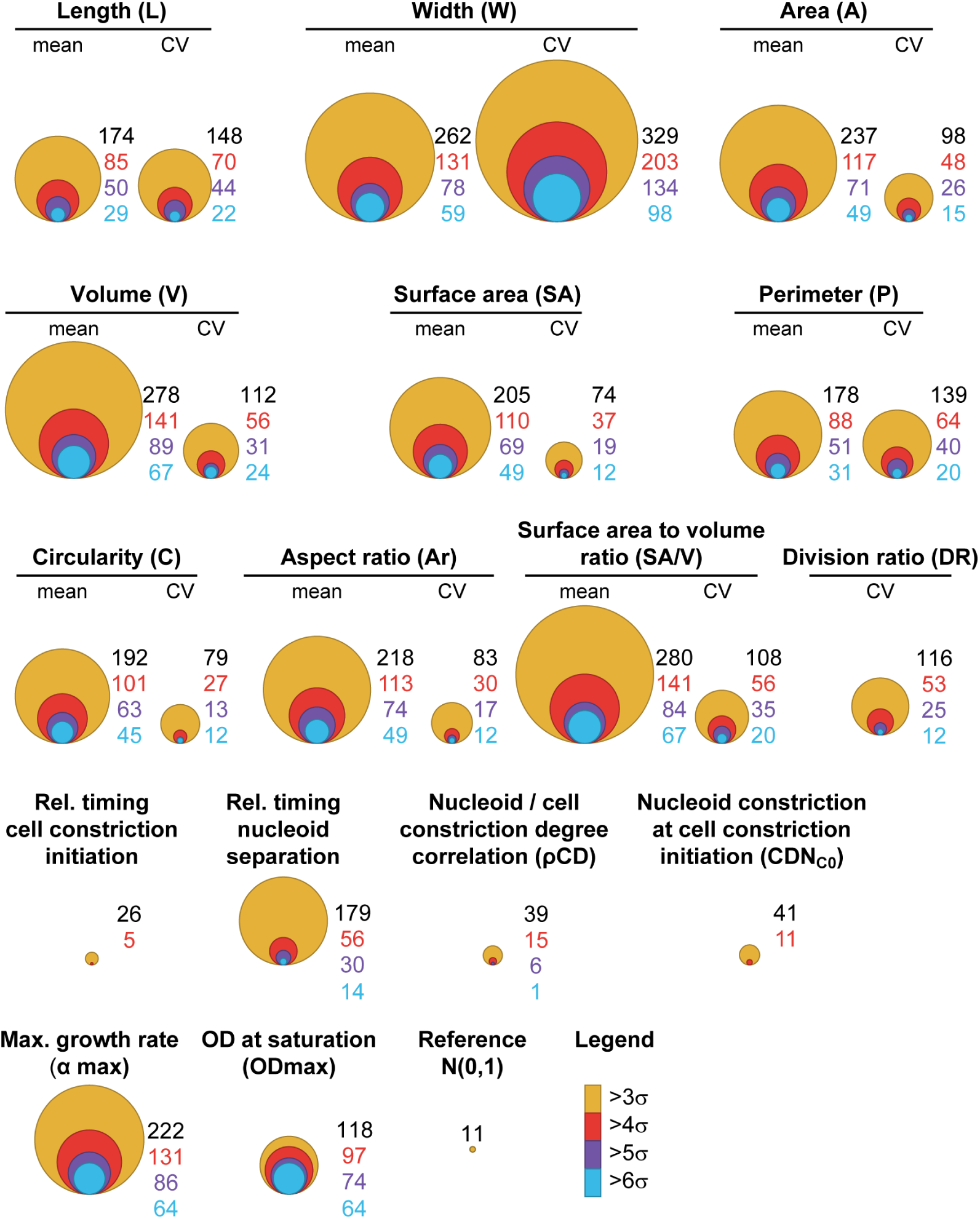
Distribution of morphological, cell cycle and growth phenotypes in the *E. coli* Keio strain collection. Bubble graphs representing, for each feature, the number of strains with a score value, *s*, beyond 3, 4, 5 or 6 times the interquartile range away from the median of the WT distribution (240 replicates), corrected by a factor 1.35 to express the deviation in terms of standard deviations (iqr ≈1.35 x *σ* for a normal distribution). The size of the circles (bubbles) reflects the number of strains with a score beyond a specific, color-coded threshold for *s*, as indicated. The ‘reference’ bubble graph illustrates the expectations from a dataset of the same size (4,227 strains), assuming a standardized normal distribution of scores (with a mean of 0 and a standard deviation of 1).

### Quantification of growth and cell cycle features across the genome

From the images, we also calculated the degree of constriction for each cell, and determined the fraction of constricting cells in the population for each strain (see Materials and methods). From the latter, we inferred the timing of initiation of cell constriction relative to the cell cycle (Collins & Richmond, 1962; Powell, 1956; Wold et al, 1994). In addition, the analysis of DAPI-stained nucleoids with the objectDetection module of Oufti (Paintdakhi et al, 2016) provided additional parameters, such as the number of nucleoids per cell and the fraction of cells with one versus two nucleoids. From the fraction of cells with two nucleoids, we estimated the relative timing of nucleoid separation (Collins & Richmond, 1962; Powell, 1956; Wold et al, 1994). We also measured the degree of nucleoid constriction in each cell for each strain and compared it to the degree of cell constriction to obtain the Pearson correlation between these two parameters, as well as the average degree of nucleoid separation at the onset of cell constriction (Appendix Fig S1H). As a result, each strain was associated with 5 cell cycle features (Dataset EV2), in addition to the 19 morphological features and 2 growth features mentioned above (see Table S1).

We found that 231 gene deletions were associated with at least one dramatically altered (|*s*| ≥ 3) cell cycle feature (Fig 2, Dataset EV2). From the growth curves, we identified over 263 mutants with severe (|*s*| ≥ 3) growth phenotypes (Fig 2, Dataset EV2) despite the growth medium being supplemented with amino acids.

### Severe defects in growth, cell morphology or the cell cycle associate with a wide variety of cellular functions

For each feature, the genes deleted in mutant strains with a |*s*| ≥ 3 encompassed a wide range of cellular functions based on a distribution analysis of Clusters of Orthologous Groups (COGs) of proteins (Fig 3 and Appendix Fig S5). This diversity highlights the high degree of integration of cell morphology and the cell cycle in overall cellular physiology.

Certain COGs were statistically enriched for some phenotypes (Fig 3). We recovered expected associations, such as category D (cell cycle control, cell division and chromosome partitioning) with high mean length (<L>) and high length variability (CV_L_) and category M (cell wall/membrane/cell wall biogenesis) with high mean width (<W>) (Fig 3A). Indeed, defects in DNA partitioning and repair can lead to a cell division block (Mulder & Woldringh, 1989), and impairment in cell envelope biogenesis has been reported to cause cell widening (Bean et al, 2009; Lee et al, 2014). COG categories associated with translation or some aspect of metabolism were, not surprisingly, enriched in mutants with growth defects (Fig 3B). Category H was enriched among small (*s* < −3) mutants. This category encompasses a number of pathways important for general aspects of metabolism (e.g., biosynthesis of panthothenate, electron carriers, biotin, and chorismate), suggesting that their impairment affects cell size in a manner similar to nutritional restriction.

**Figure 3.**
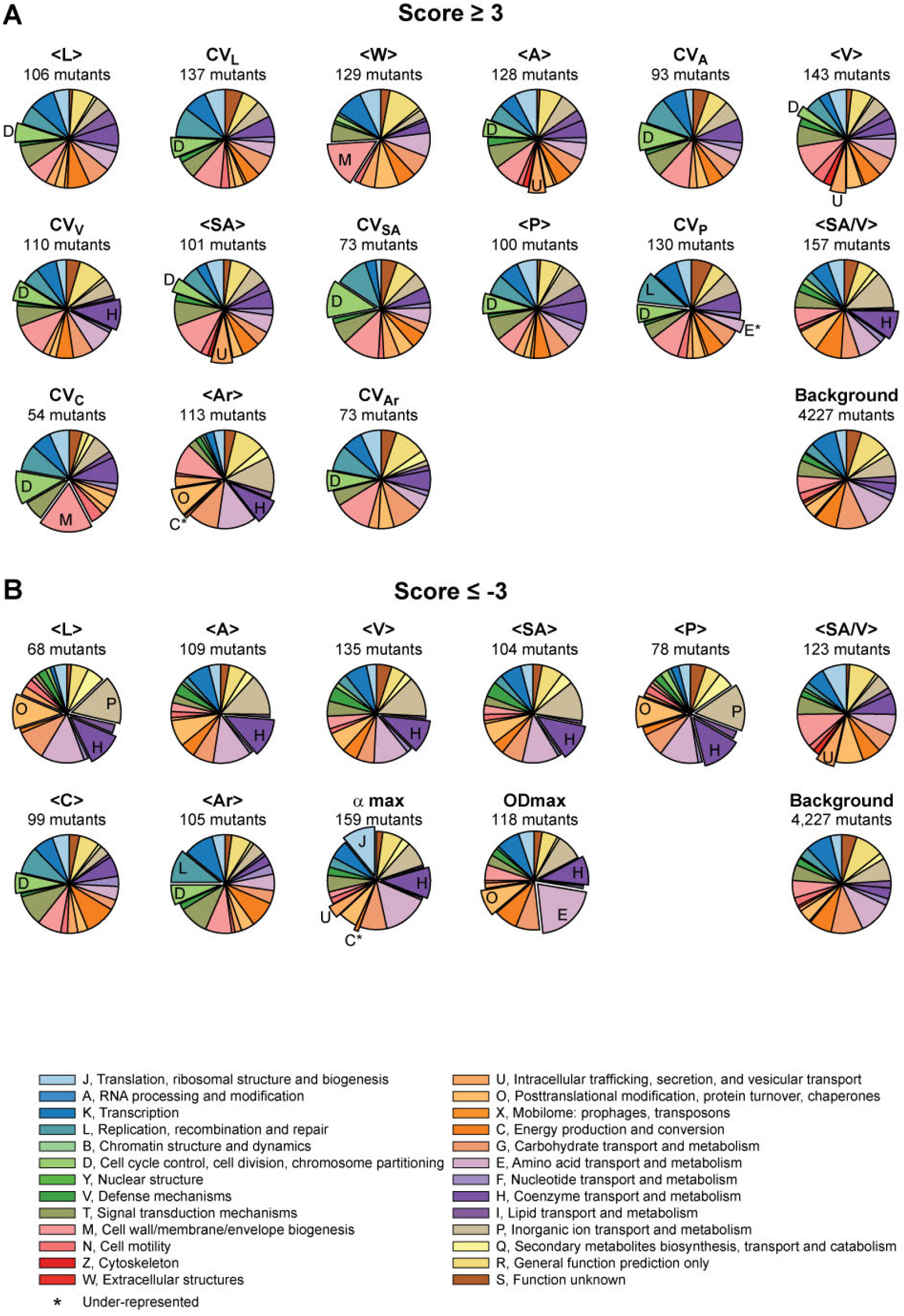
Feature-based COG enrichment analysis. **A.** Pie charts representing, on a feature-by-feature basis, the relative distribution of COG categories among the gene deletion strains associated with *s* ≥ 3. The enriched COG categories are labeled and highlighted with an exploded pie sector. The under-represented COG categories are further highlighted by an asterisk. Enrichments and under-representations with an associated (FDR-corrected) q-value < 0.05 were considered significant. Only morphological and growth features with at least one enriched or under-represented COG category are represented. **B.** Same as A but for *s* ≤ −3.

Often, these COG enrichments were carried over to features (area, volume, perimeter, circularity, etc.) that directly relate to width and length (Fig 3). However, we also observed differential COG enrichments even for highly related features, highlighting the importance of considering features beyond mean and CV of length and width. For example, category U (intracellular trafficking, secretion and vesicular transport) was enriched among mutant strains with high mean area (<A>) and volume (<V>) but normal <L> or <W> (Fig 3A), suggesting that small deviations in length and width can combine to produce significant differences in area and volume. On the other hand, deletions in category C genes (energy production and conversion) were normally represented for most phenotypes, but were conspicuously underrepresented among mutants with high mean shape factors, suggesting that deletion of these genes was barely associated with a high aspect ratio (<Ar>) (Fig 3). Thus, deletion of genes involved in energy and conversion can influence the size of the cell without affecting its shape (aspect ratio and circularity), implying that a defect in length is often accompanied by a defect in width in this category of mutants.

### High-dimensional classification of the morphological mutants

While the gene deletion annotation of the Keio library is not perfect, our large dataset provided a powerful platform to examine global trends and to identify gene function enrichments in phenotypic classes of mutants with |*s*| ≥ 3. First, we considered morphological phenotypes. Instead of ranking strains on a feature-by-feature basis, we sought to classify strains based on their combination of features, or ‘phenoprints’, to better capture the phenotypic complexity of morphology. In addition to the 19 morphological features, we included the two growth-related features (OD_max_ and *α*_max_) in the phenoprint because growth rate is often assumed to affect cell size. This assumption stems from the early observation that bacterial cell size (mean cell mass) scales with growth rate when the latter is modulated by varying the composition of the culture medium (Schaechter et al, 1958). This scaling relationship has historically been referred to as the ‘growth law’.

The combination of the 21 scores was used to classify a dataset composed of 240 wild-type replicates (controls) and 985 mutant strains with a |*s*| ≥ 3 for at least one morphological or growth feature. A principal component analysis (PCA) of this dataset was not useful, as it identifies strong outliers with severe phenotypes but failed to separate most strains (Appendix Fig S6). Therefore, to identify strains with similar phenoprints, we turned to the machine learning ‘t-distributed stochastic neighbor embedding’ (tSNE) algorithm (van der Maaten & Hinton, 2008). Unlike PCA, which identifies the principal components that most explain the variance in a dataset, the principle of tSNE is to minimize distances between datapoints with high mutual information. Thus, tSNE can be used to emphasize similarities, rather than dissimilarities. Taking advantage of the stochastic nature of tSNE, we generated 100 independent maps and used the density-based clustering algorithm dbscan (Ester et al, 1996) to identify strains that reproducibly (>90% of the time) clustered together in the tSNE maps. Using this combined tSNE-dbscan, we found that the wild-type replicates clustered together to form the ‘WT’ island while 90 of the mutant strains consistently separated in 22 islands (Fig 4A).

**Figure 4.**
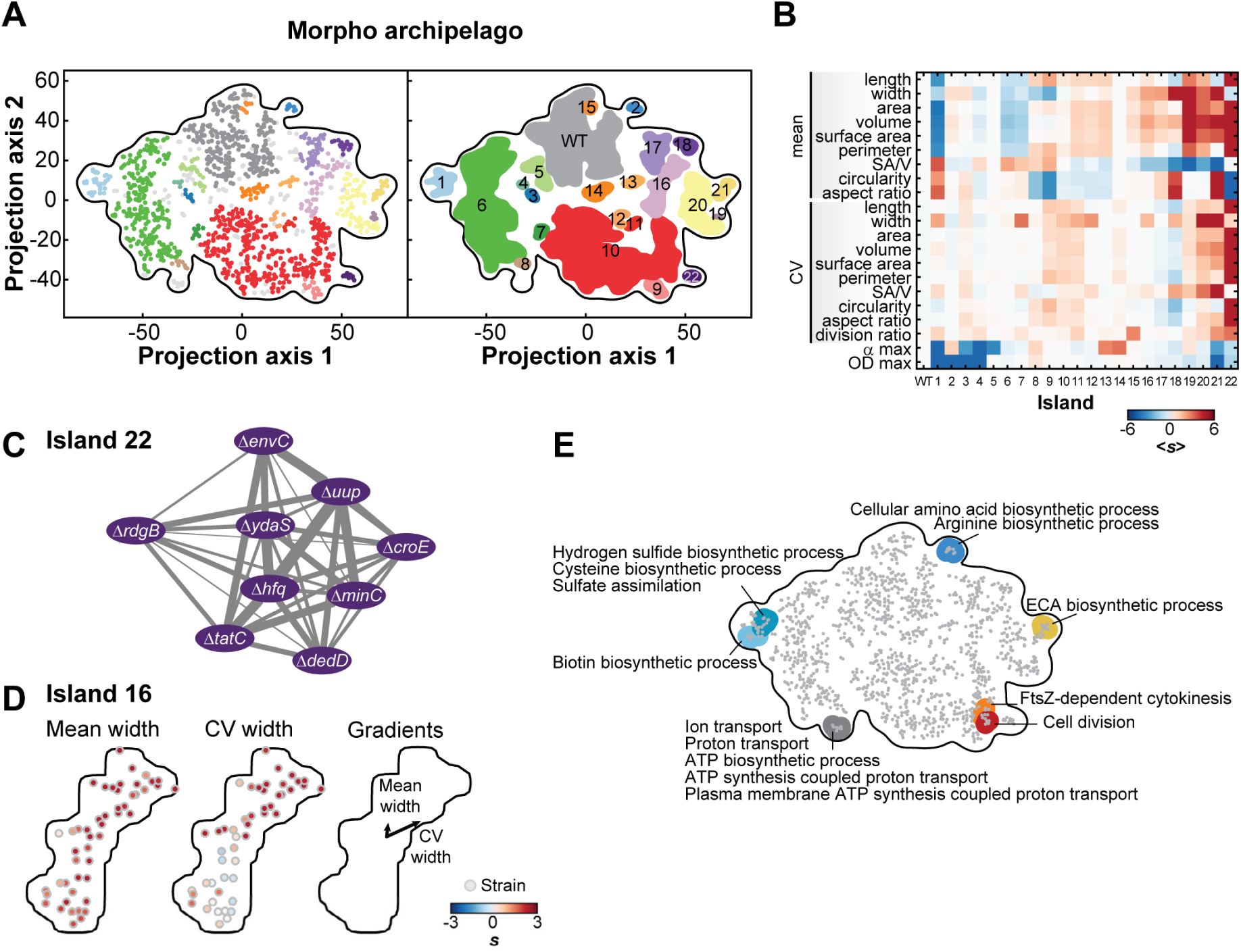
The morpho archipelago. **A.** Average 2D tSNE map of the 985 strains with at least one morphological or growth feature with a |*s*| > 3, plus the 240 independent WT replicates used as controls. Color-coded islands resulting from the dbscan algorithm (*ϕ* = 4.9, minPoints = 3) were defined by groups of strains clustering together with the same dbscan parameters in more than 90% of the generated tSNE maps. Dots in the scatter plot (left graph) represent strains color-coded based on their island affiliation (right graph). Light grey dots represent strains that were not consistently (less than 90% of the time) associated with one of the islands. **B.** Heatmap showing, for each island, the average score of each morphological and growth feature used for the construction of the map. **C.** Network representation of island 22 grouping filamentous mutants. The weights were directly derived from the average distances between corresponding mutant strains in the 2D tSNE maps. **D.** Internal structure of the mean and CV of width in island 16. The average gradients of phenotypes over the area of the island 16 are represented with arrows. **E.** Shown are the enriched “biological process” GO terms associated with false discovery rate below 0.05. For a list of all enriched GO terms, see Table S2.

Each island of the “morpho archipelago” was characterized by an average phenoprint (Fig 4B), with a given feature often segregating in different islands. For example, slowly growing (low *α*_max_) mutants were found in both islands 1 and 5, but mutants in island 1 were, on average, small whereas mutants in island 5 were morphologically like WT (Fig 4B). Thus, island 5 illustrates a group of strains that departs from the growth law, as they produce cells that are as large as WT despite growing slower.

### Genes, functions and pathways associated with cell size and shape

Our tSNE classification identified many new genes associated with specific phenotypes, even extreme ones. For example, island 22 grouped strains characterized by cells that were very long and highly variable in length (and therefore area, volume, surface area, perimeter, and aspect ratio), but had a comparatively normal width (Fig 4B). Such a cell filamentation phenotype has been well studied, and our classification recovers expected gene deletions such as ∆*minC*, ∆*envC*, ∆*tatC*, ∆*hfq* and ∆*dedD* (Fig 4C and Fig EV1A) (Adler et al, 1967; Gerding et al, 2009; Rodolakis et al, 1973; Stanley et al, 2001; Tsui et al, 1994). Island 22 also includes 4 gene deletions (∆*uup*, ∆*rdgB,* ∆*croE* and ∆*ydaS*) that were unknown for their cell filamentation phenotype (Fig 4C and Fig EV1A), suggesting new or unappreciated functions connected to cell division. For example, Uup is a DNA-related protein known to prevent the precise excision of transposons (Hopkins et al, 1983). The working model postulates that Uup interacts with the replisome to prevent replication forks stalling at the repeated sequences flanking transposons, a step required for the formation of a Holliday junction and excision (Murat et al, 2006). Replisomes also frequently stop at other chromosomal regions during replication, which can cause DNA lesions (Cox et al, 2000). If DNA damages are left uncorrected, they lead to inhibition of cell division. The cell filamentation phenotype associated with the deletion of *uup* may suggest that Uup plays a fundamental role in limiting replisome from stalling under normal growth conditions, possibly at structured DNA sites such as inverted repeats.

The ∆*rdgB* mutant suggests another underappreciated aspect of cell division (Fig 4C and Fig EV1A). RdgB is an enzyme that reduces the levels of non-canonical purines deoxyinosine (dITP) and deoxyxanthosine (dXTP) to prevent DNA damage associated with their incorporation into the chromosome; *rdgB* is essential for viability in a *recA*^*−*^ background (Budke & Kuzminov, 2010; Lukas & Kuzminov, 2006). The high frequency of cell filamentation among ∆*rdgB* cells, despite the presence of functional recombination machinery, underscores the importance of a tight control of dITP and dXTP levels in the cell.

The two remaining mutants in island 22 were strains deleted for the cryptic prophage genes *croE* and *ydaS* (Fig 4C and Fig EV1). They illustrate how this screen can identify functions for genes that are not expressed under normal growth conditions. Genes in the Keio collection were deleted by an in-frame replacement of a kanamycin resistance cassette that has a constitutive promoter and no transcriptional terminator to ensure expression of downstream genes in operons (Baba et al, 2006). However, for repressed or poorly expressed operons, the kanamycin cassette promoter can lead to unregulated expression of downstream genes in operons. This was the case for the *croE* and *ydaS* deletion strains, as cells became normal in length when the kanamycin cassette was excised (Fig EV1B and C). These results, together with the normal CV_L_ and <L> scores associated with the deletions of the downstream genes (Fig EV1B and C, Dataset EV2), suggest that it was not the loss of *croE* and *ydaS* but rather the expression of the prophage genes located directly downstream (*ymfL* and *ydaT*, respectively) that was responsible for the observed cell filamentation phenotype. Consistent with our hypothesis, it has been postulated that *ymfL* is involved in cell division (Burke et al, 2013; Mehta et al, 2004; Wang et al, 2010). While *ymfL* probably encodes a cell division inhibitor, the prophage gene *ydaT* likely inhibits cell division indirectly by acting on DNA replication or segregation, given the absence of well-segregated DAPI-stained nucleoids in filamentous ∆*ydaS* cells still carrying the kanamycin cassette (Fig EV1C).

Note that each island represented a continuum of phenotypes dominated by the features that lead to their clustering into one common island. Beyond the global segmentation of the morpho-space, each island displayed some internal structure. This is illustrated in Fig 4D, which shows the gradient of the dominating (<W>) and secondary (CVw) features within island 16. This fine internal organization reflects the objective function of the tSNE algorithm, which seeks to minimize distances between similar phenoprints. This property provided us with an excellent layout to consider tSNE maps as networks (e.g., Fig 4C), from which we could perform local functional enrichment analyses based on gene ontology (GO) terms. This approach enabled the functional annotation of the tSNE networks while taking into account the map topology without explicit clustering (see Materials and methods). This analysis highlighted both expected and surprising functional associations with specific morphological phenoprints (Fig 4E). For example, the phenoprint dominated by a small cell size and growth defects (island 1), which is a hallmark of starved cells, was not surprisingly associated with an enrichment of strains deleted for genes involved in sulfate assimilation and biotin metabolism (Fig 4E).

Another example is the enrichment of genes in the Enterobacterial Common Antigen (ECA) biosynthesis pathway (Fig 4E) among gene deletions that dramatically affected cell width control (island 21). ECA mutants tended to be wider and often lost their rod shape to form rounder cells, as shown by their high aspect ratio score (Fig EV2A, Dataset EV2). This phenotype is reminiscent of the cell shape defects caused by drugs (e.g., fosfomycin) that inhibit peptidoglycan synthesis (Kahan et al, 1974; Marquardt et al, 1994). Island 21 included other cell envelope mutants with a similar phenotype, such as gene deletions related to colanic acid (CA) biosynthesis (Dataset EV2). These results are consistent with recent studies showing that misregulation of cell shape can be caused by a competition between the ECA, CA and peptidoglycan precursor pathways for the same undecaprenyl phosphate lipid carrier (Jorgenson et al, 2016; Jorgenson & Young, 2016). The cell-width phenotype of gene deletions in the neighboring island 20 could be rationalized with a similar competition argument, as several of them are related to central metabolism (Dataset EV2). The metabolic genes may be essential for the production of key metabolites important for the synthesis of cell envelope precursors. The ∆*rapZ* strain, which had a severe cell width phenotype (Fig EV2B, island 20), may be an example. RapZ post-transcriptionally regulates the amount of GlmS (Gopel et al, 2013), which catalyzes the first committed step away from the upper glycolysis pathway and toward the synthesis of a central precursor (UDP-N-acetyl-*α*-D-glucosamine) for the biogenesis of peptidoglycan and ECA.

We also identified pathways associated with phenotypes that were not easy to rationalize. Deletion of genes encoding the high-affinity phosphate transporter subunits (PstA, PstC and PstS), the associated histidine kinase (PhoR) and the adaptor protein (PhoU) led to a thin phenotype (Fig EV2C), without significantly slowing down growth (Dataset EV2). The absence of a growth defect is expected, as the growth medium is rich in phosphate that can be taken up by the low-affinity phosphate transporters. Thus, the cell width reduction cannot be associated with phosphate starvation.

Deletion of several genes encoding subunits of ATP synthase, which results in a metabolic switch to fermentation, led to a decrease in average cell width (Fig EV2C). Cultures of these deletion strains did not grow slower (*s* > 0 for *α*_max_) than WT (Dataset EV2). Furthermore, they were imaged at an OD_imaging_ at least 3 times smaller than their OD_max_, indicating the cell width phenotype could not be linked to the inability of some of these strains to grow to high cell density (*s* < −3 for OD_max_). This result suggests that either the ATP synthase itself or differences in metabolism alter cell shape and size independently of growth rate.

Another surprise was the lack of clustering, and therefore the absence of association, among mutants expected to affect fatty acid metabolism; instead, fatty acid mutants displayed a variety of phenotypes (Dataset EV2). Previous work has shown that a reduction of fatty acid synthesis through drug treatment or deletion of the fatty acid biosynthetic gene *fabH* results in a thinner and shorter cell phenotype (Yao et al, 2012). Conversely, an excess of fatty acids through overexpression of the regulator *fadR* or by addition of exogenous fatty acids leads to wider and longer cells (Vadia et al, 2017). These results have led to a simple model in which the amount of fatty acids and, by extension, the level of lipid synthesis determine cell size. Our data suggest a potentially more complex relationship between phospholipids and cell morphology. This is illustrated by the ∆*fadR* and ∆*fabF* strains, which were thinner (s < −8), but also longer (s > 3.5), than the parental strain (Fig EV2D). Remarkably, the width and length defects were compensatory such that the ∆*fadR* and ∆*fabF* mutants retained a normal cell area (Fig EV2D). FadR is a bifunctional transcriptional factor that activates fatty acid synthesis and represses *β*-oxidation, while FabF is a fatty acid chain elongation enzyme. Based on their metabolic profiles, both ∆*fadR* and ∆*fabF* strains have significant changes in the levels of phospholipids containing saturated versus unsaturated acyl chains (Fig EV2D) (Fuhrer et al, 2017; Garwin et al, 1980; Nunn et al, 1983). These results suggest that an altered phospholipid composition, such as changes in the degree of fatty acid saturation, may be another important factor that determines the dimensions of the cell.

### Identification of genes affecting nucleoid separation and cell constriction dynamics

We applied the same tSNE analysis to the seven cell cycle and growth features of the 397 strains displaying a severe defect (|*s*| ≥ 3) for at least one cell cycle or growth feature. The 240 independent wild-type replicates were included in the analysis as controls. We robustly identified a WT island and 12 distinct mutant islands in this cell-cycle space (Fig 5A). Each island was characterized by an average phenoprint (Fig 5B). Islands 6 and 7 were phenotypically close to WT. Islands 3 and 10 grouped mutants with growth defects and little to no cell cycle phenotypes (Fig 5B and C). The neighboring islands 1, 2, 9 and 11 were dominated by cell growth features with some combination of nucleoid separation and cell constriction defects. Three islands—islands 4, 8 and 12—grouped interesting gene deletion strains with cell cycle phenotypes and no significant growth defects (Fig 5B and C).

**Figure 5.**
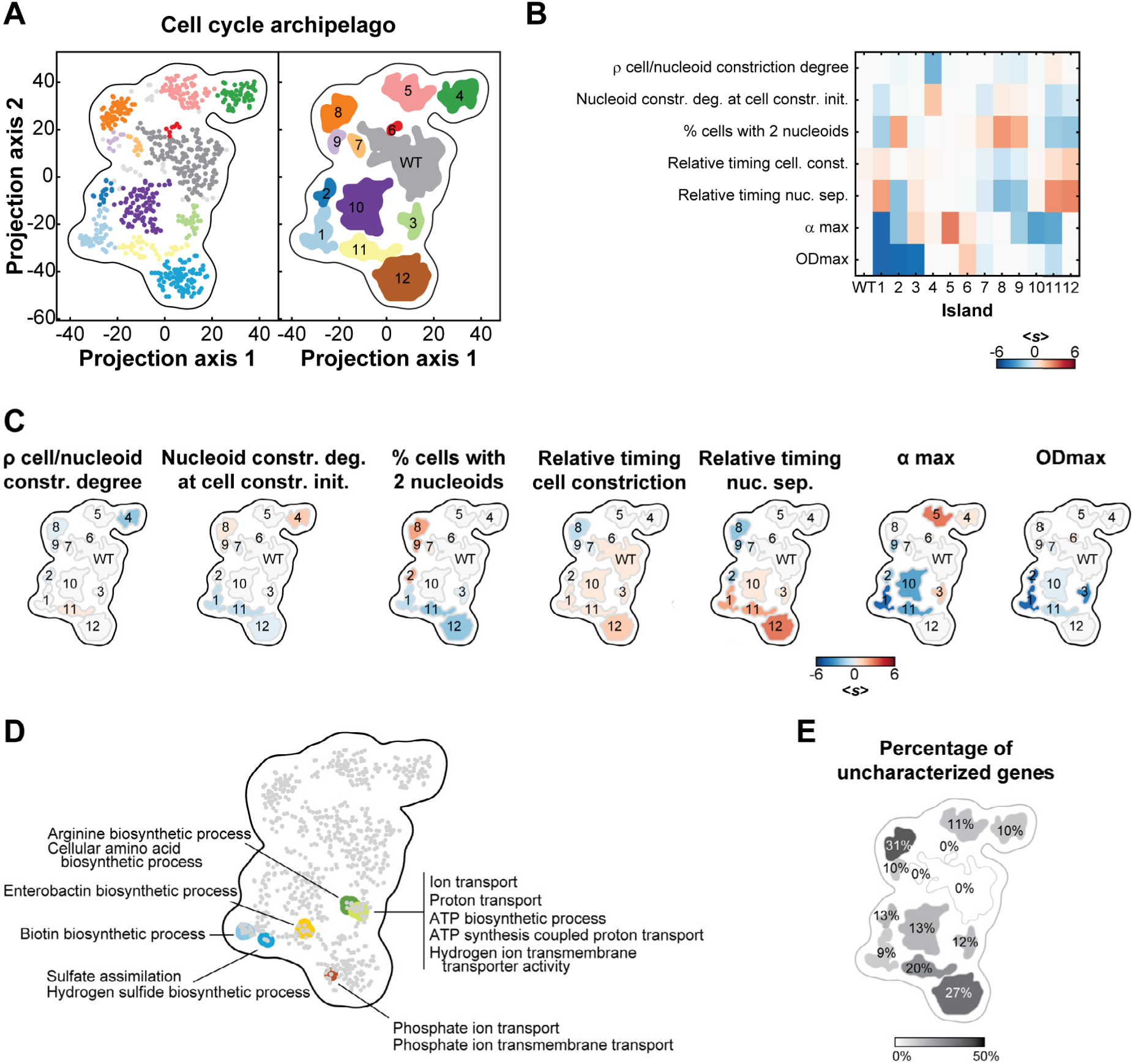
The cell cycle archipelago. **A.** Stable islands in the cell cycle archipelago. The cell cycle and growth phenoprints were used to map the 397 mutant strains with at least one cell cycle or growth feature with a |*s*| ≥ 3, as well as the 240 independent WT replicates, in 2D using tSNE. As for the morpho archipelago, the data were clustered using the dbscan algorithm. The groups of strains clustering (dbscan parameters *ϕ* = 4.28, minPoints = 3) together in more than 90% of the maps defined an island. Dots in the scatter plot on the left represent strains and are colored with the same color code as for the island on the right graph. Light grey dots represent strains that were not consistently (less than 90% of the time) associated with one of the island. **B.** Heatmap showing, for each island, the average score of each cell cycle and growth feature used for the construction of the map. **C.** Islands shown in panel A were colored according to their average score for each listed feature. **D.** Shown are the enriched “biological process” GO terms associated with false discovery rate below 0.05. For a list of all enriched GO terms, see Table S2. **E**. The islands in the cell cycle archipelago were colored according to the proportion of ‘y-genes’ (genes of unknown function).

Functional analysis on all strains identified GO term enrichments with phenoprints that show strong growth phenotypes (Fig 5D). We did not find any GO term enrichment associated with cell cycle phenotypes independently of growth defects. Furthermore, the proportion of genes of unknown function (so-called ‘y-genes’) was the highest (~ 30%) for cell cycle-specific islands 8 and 12 (Fig 5E). These observations highlight the limited extent of our knowledge about the genetic basis of nucleoid and cell constriction dynamics, compared to cell growth.

Our analysis of nucleoid separation and cell constriction provided a genome-wide perspective on the processes affecting DNA segregation and cell division. While each event has been investigated for years at the molecular level, we know little about their coordination. We found that nucleoid separation is tightly correlated with the initiation of cell constriction across the ~ 4,000 deletion strains (*ρ* = 0.65, 95% confidence interval [0.63, 0.66], Fig 6A) and at the single-cell level (Appendix Fig S1H). A well-known genetic factor involved in this coordination is MatP (Mercier et al, 2008). This DNA-binding protein organizes and connects the chromosomal terminal macrodomain (*ter*) to the division machinery (Espeli et al, 2012). Consistent with this function, we observed that the ∆*matP* mutant, which segregated into island 8, failed to coordinate nucleoid separation with cell constriction and separated its nucleoid early while dividing at about the same cell age as WT (Fig 6A-C). The early separation of nucleoids is in agreement with the early segregation of sister loci within the *ter* region in the ∆*matP* mutant (Espeli et al, 2012; Mercier et al, 2008) and with the proposed role of MapP in linking DNA segregation to cell division (Mannik and Bailey. 2015). The remaining 47 strains from island 8, which also displayed an early nucleoid separation phenotype (Fig 6B), were deleted for genes that had either uncharacterized functions or functions unrelated to nucleoid dynamics (Dataset EV2). Deletion mutants that clustered closely to the ∆*matP* mutant within island 8 displayed a WT-like cell constriction profile as well (Fig 6B). Examples include mutants that lack the lysophospholipase L2, PldB, the DNA repair polymerase, PolA, or the poorly characterized protein, YfjK (Fig 6B, D). YfjK is a predicted helicase that has been associated with sensitivity to ionizing radiations (Byrne et al, 2014). On the other side of the island, mutants were characterized by not only an early nucleoid separation, but also early initiation of cell constriction (Fig 6A, B and E), to the point that the timing of cell constriction and nucleoid separation was virtually the same. This is illustrated with ∆*ycjV* and ∆*hlsU* (Fig 6E). YcjV is a predicted ABC transporter ATPase. HslU has two functions in the cell, one as a subunit in a protease complex with HslV, and the other as a chaperone (Seong et al, 2000; Slominska et al, 2003). Since we did not observe any significant defect in cell constriction timing for the ∆*hslV* mutant, the ∆*hlsU* phenotype is more likely linked to the chaperone activity.

**Figure 6.**
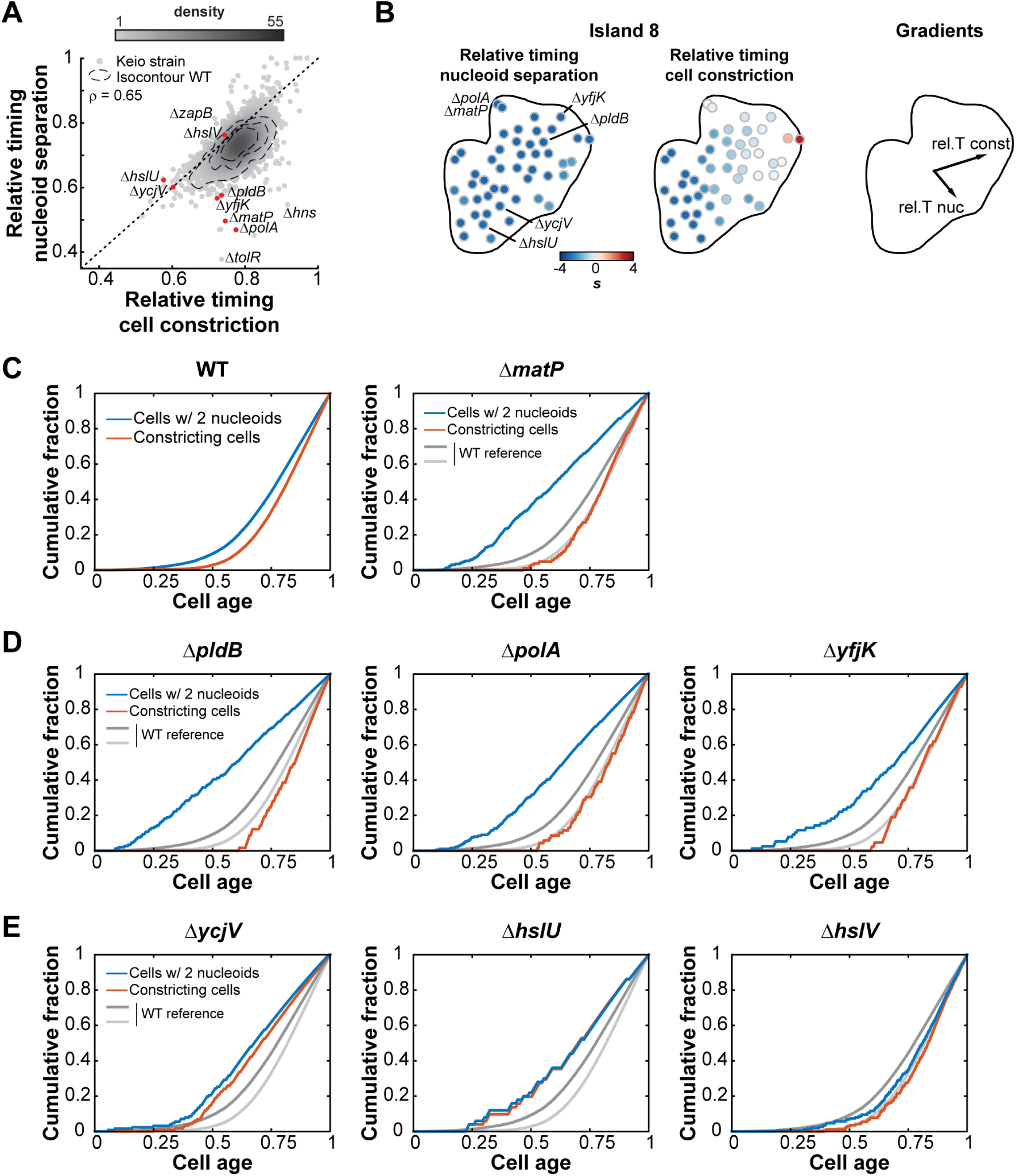
Nucleoid separation and cell constriction dynamics. **A.** Scatter plot of the relative timing of cell constriction versus the relative timing of nucleoid separation. The gray scale indicates the density of dots in a given area of the chart. The dotted contours represent the 0.5, 0.75 and 0.95 probability contours of the 240 WT replicates. The Pearson correlation is *ρ* = 0.65, with a 95% CI = [0.63, 0.66]. The black dotted diagonal represents the line where a strain should be if both nucleoid separation and cell constriction happens at the same time. Red dots highlight strains shown in panels **C**, **D** and **E**. **B.** Close-up view of the cell cycle island 8 with each dot representing a Keio strain colored according to the two features driving the clustering of the 48 strains. The relative timing of nucleoid separation is the dominant feature of island 8, while the relative timing of cell constriction drives the layout of the strains within the island. **C.** Average dynamics of nucleoid separation and cell constriction for WT and the ∆*matP* mutant strain. The cumulative distributions of the fraction of cells with two nucleoids (blue) and of the fraction of cells with a constriction degree above 0.15 (red) were plotted against cell age. Cell age was calculated according to the rank of each cell based on their cell length with the formula 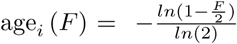, where F represents the fraction of cells with a cell length equal or below the length of cell *i* (Wold et al, 1994). **D.** Same plots as in **C** for three strains clustering in island 8 with the ∆*matP* strain. The WT curves shown in **C** were plotted in gray for comparison. **E.** Same plots as in **C** for two island-8 strains, ∆*ycjV* and ∆*hslU*, and the ∆*hslV* strain, which does not partition in island 8.

### Identification of cell size control mutants

How cells achieve size homeostasis has been a longstanding question in biology. While the control mechanism at play remains under debate (Amir, 2014; Campos et al, 2014; Harris & Theriot, 2016; Ho & Amir, 2015; Iyer-Biswas et al, 2014; Taheri-Araghi, 2015; Taheri-Araghi et al, 2015; Tanouchi et al, 2015; Wallden et al, 2016), we and others have shown that under the growth conditions considered in this study, *E. coli* follows an adder principle in which cells grow a constant length (∆L) before dividing (Campos et al, 2014; Taheri-Araghi et al, 2015). We sought to use this screen to survey the role of genes in cell length control. We first explored the relationship between mean length (<L>) and length variability (CV_L_) among mutants. On average, short mutants (*s* ≤ −3, n = 68) had a normal CV_L_ (p-value = 0.98), while long mutants (*s* ≥ 3, n = 106) displayed a greater CV_L_ (p-value = 8.18 10^−11^, Fig 7A).

**Figure 7.**
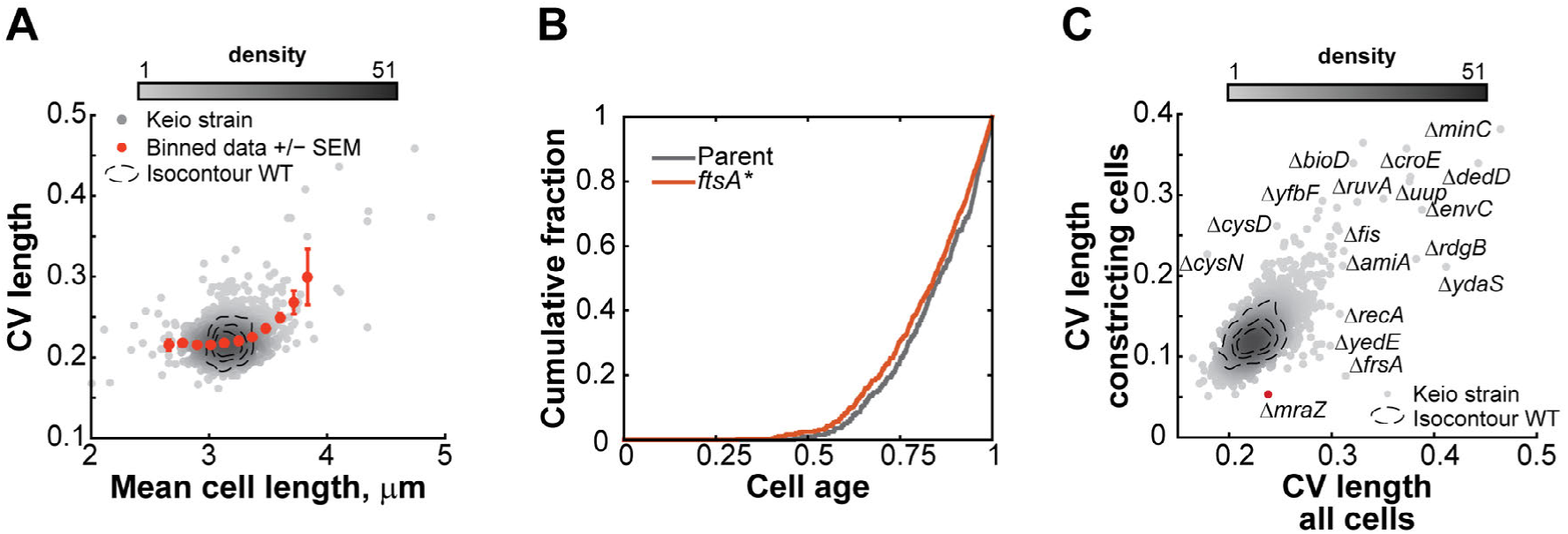
Cell length regulation mutants. **A.** Scatter plot of the mean cell length versus the CV of the length for all the strains. The gray color levels indicate the density of points in the vicinity of each strain. The orange dots and error bars represent the mean and standard error of the mean per bin. **B.** The cumulative distribution of the proportion of constricting cells for the *ftsA\** mutant and its parent were plotted against cell age. We measured the degree of constriction of all cells using Oufti. For each strain, we had two independently acquired image sets (n = 496 and 1,168 cells for the WT replicates and n = 692 and 1,506 cells for the *ftsA*\* replicates). The distributions of the degree of constriction were not significantly different among the four datasets at a threshold p-value of 0.01 using a Kruskal-Wallis multi-comparison test. Moreover, Bonferroni-corrected post-hoc pairwise tests did not allow a distinction between WT and *ftsA*\* samples. **C.** Scatter plot of the CV of the cell length for the whole population versus the CV of the cell length for constricting cells only. The contour lines represent the 0.5, 075 and 0.95 probability envelopes of the 240 independent WT replicates. The gray color levels indicate the density of points in the vicinity of each strain. The ∆*mraZ* strain discussed in the text is highlighted in red.

The observation that short mutants displayed, on average, a normal CV_L_ indicates that most of them regulate their length distribution as precisely as WT. These results suggest that the adder principle, and therefore the relative timing of cell division, is just as precise in short mutants as in WT cells. This result is interesting because short mutants have traditionally received a lot of attention in cell size control studies. A well-known short mutant in *E. coli* is the *ftsA\** strain (WM1659), which is thought to misregulate size control by triggering division prematurely (Geissler et al, 2007; Hill et al, 2012). However, when we imaged the *ftsA\** mutant (n = 2,198 WM1659 cells), we found that, similar to the trend shown by short mutants in our screen, *ftsA\** cells constrict at a similar cell age as WT (Fig 7B). In hindsight, this result makes sense since the WT and *ftsA*\* strains have a similar doubling time (72.3 ± 2.2 min versus 69.0 ± 3.2 min, mean ± standard deviation, n = 4), consistent with Geissler et al (2007) and therefore take the same amount of time to divide. Perhaps a more appropriate way to consider short mutants with normal CV_L_ is not as mutants that have a premature division, but as ‘small-adder’ mutants since they add an abnormally small cell length increment ∆L between divisions.

Long mutants, on the other hand, tended to lose their ability to maintain a narrow size distribution, as CV_L_ increased with <L> (Fig 7A). The origin for an increase in CV_L_ may signify a loss of precision in the timing of division, but it may alternatively originate from an aberrant positioning of the division site (or both). The ∆*minC* mutant is an example of aberrantly large CV_L_ (Fig 7C) due to the mispositioning of the division site, and not due to a defective adder (Campos et al, 2014). This class of mutants can easily be identified in our dataset by their large variability in division ratios (CV_DR_). Conversely, a high CV_L_ associated with a normal variability in division ratios points to a mutant that has a more variable ∆L between divisions.

We suspected that interesting cell size control mutants might be missed by only considering CV_L_. The distribution of cell lengths in a population is a convolution of cell length distributions at specific cell cycle stages. Since there is significant overlap in length distributions between cell cycle stages, a substantial change in CV_L_ at a specific cell cycle stage (e.g., cell constriction) does not necessarily translate into obvious changes in CV_L_ of the whole population, as shown in simulations (Fig EV3). Our screen allowed us to identify constricting cells and hence to determine the cell length variability for the stage of cell constriction. This cell cycle stage-specific analysis identified ∆*mraZ* as a potential gain-of-function cell size homeostasis mutant (Fig 7C). For this mutant, division (CV_DR_, Dataset EV2) and growth rate (Eraso et al, 2014) were normal, but the length distribution of its constricted cells (CV_L_= 0.05) was remarkably narrower than that of WT constricted cells (CV_L_ = 0.12). MraZ is a highly conserved transcriptional regulator that downregulates the expression of the *dcw* cluster (Eraso et al, 2014), which includes cell wall synthesis and cell division genes (Ayala et al, 1994). Our data suggests that MraZ and the regulation of the *dcw* cluster affect the balance between cell growth and division.

### Dependencies between cellular dimensions and cell cycle progression

A fundamental question in biology is how cells integrate cellular processes. A common approach to address this question is to look at co-variation between processes or phenotypes following a perturbation (e.g., mutation, drug treatment). However, perturbations that affect the same system (i.e., perturbing a single process or pathway) can lead to misinterpretation, as the perturbation may abolish a given dependency between two features or may affect the co-varying phenotypes independently. Increasing the number of independent perturbations has two major effects. First, it alleviates the interpretation problem by averaging out the specific effect associated with each perturbation (Collinet et al, 2010; Liberali et al, 2015; Sachs et al, 2005). The Keio collection consists of mutants affected in a wide variety of cellular processes, allowing us to examine whether specific features correlate across many different genetic perturbations. Second, the large number of genetic perturbations increases our confidence in any calculated correlation (or lack thereof) between features. To illustrate, let’s consider two independent features (i.e., true correlation = 0). Using an analytical solution (Fisher, 1915; Fisher, 1921), one can show that the calculated 95% confidence interval (CI) for the Pearson correlation is between −0.63 and +0.63 if the sample size is 10. The calculated 95% CI is very wide because two uncorrelated features can easily appear positively or negatively correlated if the sample size is small. The 95% CI shrinks down to [-0.06, +0.06] if the sample size is 1000. Therefore, the large number and variety of mutants in our study provided an opportunity to identify global effects and dependencies between morphological, cell cycle and growth phenotypes through correlation analysis.

To build an interaction network, we used the information-theoretic algorithm ARACNE (Margolin et al, 2006). This method considers all pairwise correlations between features at the same time and identifies the most relevant connections by removing those that are weak or that can be explained via more correlated paths. In this analysis, we focused on 10 quantitative non-collinear features that describe cellular dimensions (<L>, <W>, <A>), nucleoid size (<NA>), population growth (*α*_max_, OD_max_), nucleoid separation (rel. T nuc), cell constriction (rel. T const) and their interdependency (*ρ*CD, CDN_C0_) (see Materials and methods). The resulting network recovered obvious connections, such as the relation of cell area with cell length and width. It also showed that growth rate features displayed virtually no connectivity to cell size or cell cycle features (Fig 8A). This was further illustrated by the close-to-zero correlations between growth rate and any of the 24 morphological and cell cycle features considered in our screen (Fig EV4A).

**Figure 8.**
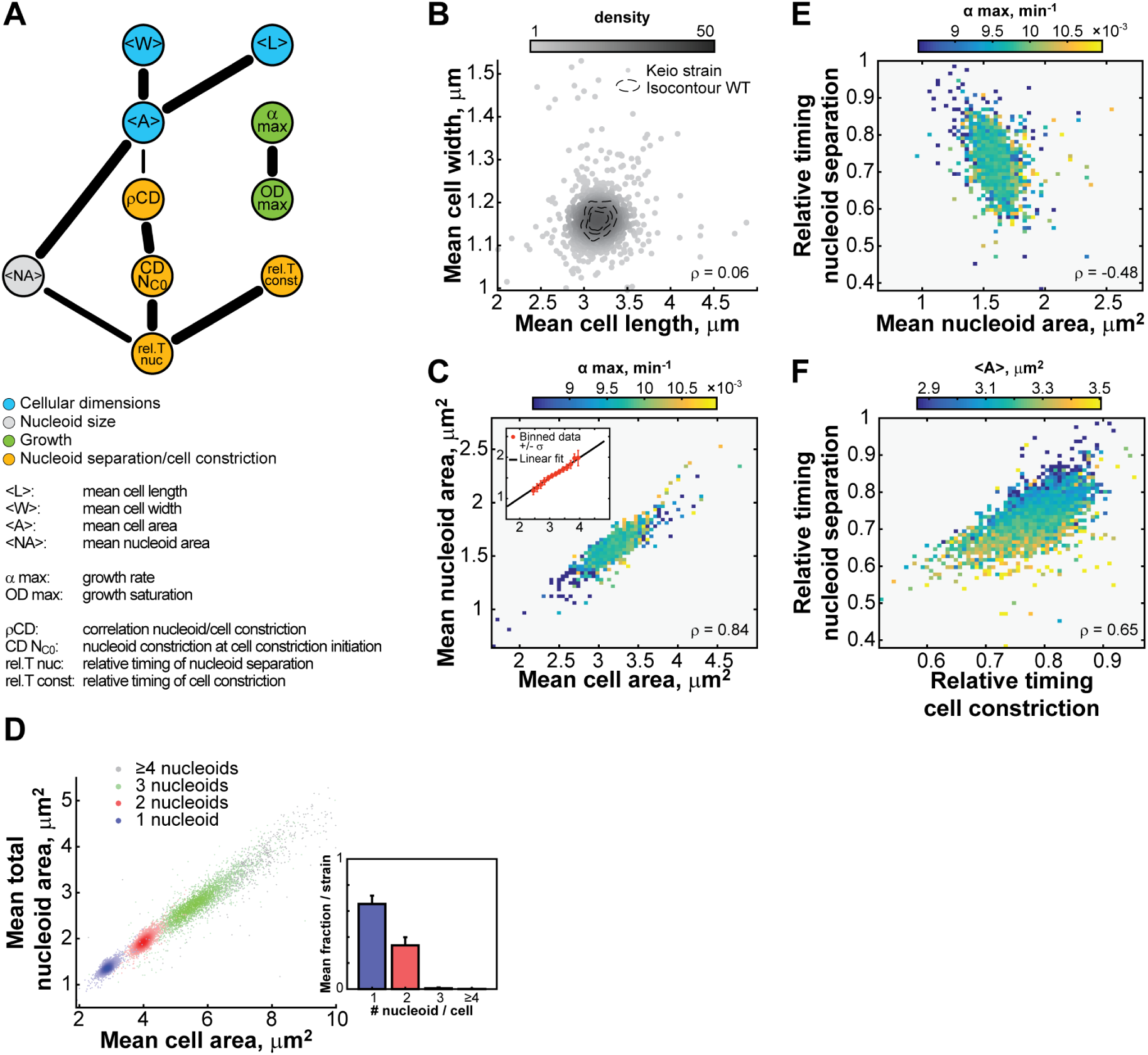
Interdependence of cell morphogenesis and cell cycle progression. **A.** Network showing the functional relationship between 10 non-collinear morphological, growth and cell cycle features. The network is an undirected network highlighting the most informative connections detected by the ARACNE algorithm. The thickness of an edge represents the fraction of the networks containing this specific edge after bootstrapping the network 200 times, from 70% (thinnest) to 100% (thickest). **B.** Scatter plot of the normalized mean cell length and mean cell width of all 4,227 Keio strains and 240 WT replicates. Each dot represents a strain, and the gray level illustrates the density of neighbors in the vicinity of each point in the graph. The dotted contours represent the 0.5, 0.75 and 0.95 probability envelopes of the 240 WT replicates. The correlation between mean cell width and mean cell length is low (*ρ* = 0.06, 95% CI [0.03, 0.09]). **C.** Heatmap showing the mean growth rate value for data binned by both mean cell area and mean nucleoid area. The cell and nucleoid areas are strongly correlated (*ρ* = 0.84, 95% CI [0.83, 0.85]). The median value of *α*_max_ per bin is color-coded according to the color scale. The inset highlights the linearity of the relationship between mean cell area and mean nucleoid area. The orange dots and error bars represent the binned data, with the black line showing the best linear fit to the binned data. **D.** Scatter plot of the mean cell area versus the mean nucleoid area for cells with 1, 2, 3 or ≥4 nucleoids for each strain. The histogram in the inset illustrates the average proportions of cells with 1, 2, 3 or ≥4 nucleoids per strain. Although there were typically few cells in each strain with 3 or ≥4 nucleoids, at least one cell with ≥3 nucleoids was detected for 61% of the strains. **E.** Heatmap showing the mean growth rate value for data binned by both the mean nucleoid area and the relative timing of nucleoid separation. The mean nucleoid area negatively correlates with the relative timing of nucleoid separation (*ρ* = −0.48, 95% CI [−0.50, −0.45]). **F.** Heatmap showing the mean cell area value for data binned by both the relative timing of cell constriction and the relative timing of nucleoid separation. The relative timing of cell constriction is strongly correlated to the relative timing of nucleoid separation (*ρ* = 0.65, 95% CI [0.63, 0.66]).

Another interesting lack of connection was between <L> and <W> (Fig 8A), as these two variables were largely uncorrelated (Fig 8B). With *n* = 4,227, our estimated Pearson correlation (*ρ* = 0.06) is associated with a narrow 95% CI between −0.02 and 0.13. By subsampling this large dataset, we can show how a decrease in sample size increases the likelihood of obtaining erroneous positive or negative correlations (Fig EV5). The lack of correlation between <L> and <W> is interesting from a cell size control standpoint. If *E. coli* was controlling its size by sensing its volume, surface area or the ratio between the two, as *Caulobacter crescentus* does (Harris et al, 2014), we would expect a global anti-correlation between length and width such that an increase in cell length would be, on average, compensated by a decrease in width, and vice versa. The lack of correlation argues that cell length and width are controlled independently in *E. coli*, at least under our growth conditions.

Some features, however, displayed strong co-variation across the 4,000 genetic perturbations. For instance, the mean cell area and mean nucleoid area (considering the sum of nucleoids in the cell) were highly positively correlated (*ρ* = 0.84, 95% CI [0.83, 0.85]), in a growth rate-independent manner (Fig 8C). In wild-type cells, nucleoid size linearly increases with cell size throughout the cell cycle (Junier et al, 2014; Paintdakhi et al, 2016). Here, we found that nucleoid size scales with cell size across ~ 4,000 mutants despite their effects on different cellular functions: small mutants had a small nucleoid size, and big mutants had a big nucleoid size (Fig 8C). This remarkable linear relationship held true regardless of the number of nucleoids per cell (Fig 8D). In addition to its strong positive correlation with the average cell size, the average nucleoid size was negatively correlated with the relative timing of nucleoid separation (*ρ* = −0.48, 95% CI [−0.50, −0.45], Fig 8E). These connections suggest a dependency between size and cell cycle features: the bigger the cell, the bigger the nucleoid is (Fig 8C) and the earlier nucleoid separation and cell constriction occur in relative cell cycle units (Fig 8E-F). This dependency is highlighted by the overall structure of the interaction network (Fig 8A), which reveals that the cell cycle features (yellow nodes) are primarily connected to the cellular dimension features (blue nodes) through the dimensions of the nucleoid (grey node).

## Discussion

In this study, we used a multi-parametric approach to quantitatively survey the role of the non-essential *E. coli* genome in cell shape, cell size, cell growth and two late cell cycle stages, nucleoid separation and cell constriction. The results provide a valuable resource of phenotypic references for both characterized and uncharacterized genes, as well as a rich dataset to explore the correlation structure between cellular dimensions, growth and cell cycle features at the system level.

The large proportion of genes and the wide variety of functions impacting cell size and shape and the progression of late cell cycle stages (Figs 2 and 3, Appendix Fig S5) underscore the degree of integration of cell morphogenesis and cell cycle progression in all aspects of *E. coli* cell physiology. It also implies that most morphological and cell cycle phenotypes cannot easily be imputed to a specific pathway or cluster of genes. In fact, genes involved in the same cellular process can have very different, and even sometimes opposing, effects. Genes associated with translation illustrate this concept. Deletion of ribosomal subunit genes leads to a diversity of morphological phenotypes, such as thin (∆*rplY*), wide (∆*rpsO*), short (∆*rpsT*), and short and thin (∆*rpsF*) (Dataset EV2). This diversity of phenotypes is also observable for deletions of genes encoding enzymes that modify ribosomal RNAs or tRNAs (e.g., ∆*rsmD* cells are long, whereas ∆*rluD* and ∆*truA* cells are wide, and ∆*mnmC* cells are long, wide and variable in size). The latter suggests an unexpected role for RNA modifications in cell morphogenesis.

Overall, this study greatly expands the number of genes associated with cell morphogenesis (874) and the cell cycle (231). Notably, it provides a phenotype for 283 genes of uncharacterized function (out of 1306 y-genes). The Keio collection, as any genome-wide deletion collection (Teng et al, 2013), is no stranger to gene duplications and compensatory mutations (Otsuka et al, 2015; Yamamoto et al, 2009). Although their occurrence can impact interpretation at the gene level, “suppressor” mutants tend to display more WT-like behavior. It is, therefore, possible that we have underestimated the actual number of mutants displaying altered phenotypic characteristics or underrated the severity of the phenotype of a given deletion. Importantly, “incorrect” mutants have no effect on our global correlation analyses, as the latter does not rely on the genetic identity of the mutants.

Our study reveals new phenotypes for previously characterized gene deletions. We have already mentioned above the unexpected filamentation phenotype of the ∆*uup* strain (Fig 4C and Fig EV1A) and have proposed a tentative connection between Uup’s known function (precise transposon excision) and DNA damage through replisome stalling. We also identified unanticipated links. For example, the requirement for lysophospholipase L2 (PldB) in the coupling of nucleoid separation and cell constriction (Fig 6D) suggests a connection between phospholipid metabolism and the coordination of late cell cycle stages. Previous works have also linked phospholipid metabolism to cell morphology (Yao et al, 2012), showing that fatty acid availability dictates the capacity of the cell envelope to expand, ultimately affecting cell size (Vadia et al, 2017). We found that mutants with an altered degree of saturated vs. unsaturated phospholipids have an abnormal length and width (Fig EV2D). It is, therefore, tempting to speculate that not only the amount, but also the composition of fatty acids plays an important role in cell shape and size control. Phospholipid composition determines the chemical and physical properties of the cell membrane (Dowhan & Bogdanov, 2002), which is known to affect the function of cell division and morphogenesis proteins such as MinD and MreB (Kawazura et al, 2017; Mileykovskaya et al, 2003).

By combining the tSNE and dbscan algorithms, we were able to cluster strains with similar phenoprints into islands (Figs 4 and 5). This granular representation of the phenotypic space allowed us to expand on well-studied archetypal phenotypes such as ‘filamentous’ and ‘fat’ (islands 22 and 21 of the morpho archipelago, respectively, see Fig 4). This classification also allowed us to populate less well-studied phenotypes, from which we can gain new insight into cell morphogenesis and the cell cycle. For example, the substantial number of thin mutants reported here may prove as valuable as fat mutants to study cell morphogenesis from a different angle. The clustering results also revealed entirely new classes of mutants. In that respect, islands 4, 8 and 12 of the cell cycle archipelago are particularly interesting because they offer a genetic toolkit to explore nucleoid and cell constriction dynamics, which have remained poorly understood despite their essential role in cellular replication.

It is important to note that the phenoprints reported in this study are tied to the specific experimental conditions of the screen. Cell size is well known to vary with nutrient conditions (Schaechter et al, 1958), indicating that the behavior of the Keio mutants may be different in other environments. Differences in growth conditions also lead to different metabolic requirements and growth limitations. For instance, none of the mutant strains auxotrophic for nucleotides were able to grow in our synthetic medium, which lacks nucleotide precursors. We also note that growth in 96-well plates likely corresponds to micro-aerophilic conditions. Accordingly, we identified morphological deviations for strains deleted for genes known to be only expressed under micro-aerophilic or anaerobic conditions, revealing new metabolic connections to cell morphogenesis. For example, deletion of *ybcF*, which is predicted to encode an enzyme involved in anaerobic purine degradation (Smith et al, 2012), results in a fat cell phenotype (Dataset EV2).

In this study, each gene deletion can be seen as a perturbation. The sheer number of perturbations (~4,000) guarantees a large number of independent perturbations and offers a unique opportunity to infer the underlying correlation structure between the different phenotypes (Fig 8A). Such relationships, or lack thereof, can be very informative. For instance, we found that growth rate is not predictive of cell size. This is an interesting finding because current theoretical models of cell size control generally include growth rate as a variable. Recently, the original growth law was modified to include a second variable, the C+D period (time of DNA replication + time between the end of DNA replication and cell division), that influence cell size (Si et al, 2017; Zheng et al, 2016). Although we do not have specific measurements of C+D periods and thus cannot directly compare our results to this general growth law, the observed lack of correlation between growth rate and cell size remains surprising, especially for the fast growing mutants (Appendix Fig S7). Growth inhibition and nutrient limitation experiments have shown that the C+D period increases with slower growth rates (Kubitschek & Newman, 1978; Si et al, 2017; Wallden et al, 2016). It is therefore possible that a lengthening of the C+D period in slow growing mutants compensates for the growth rate difference, masking the relationship between the growth rate and cell size in our dataset. However, under our nutrient-growth conditions of overlapping DNA replication cycles (Appendix Fig S1A), the C+D period is supposed to have reached a plateau, i.e., a minimal value that remains constant at even faster growth rates (Cooper & Helmstetter, 1968; Helmstetter, 1968; Si et al, 2017; Wallden et al, 2016). Therefore, we would not expect fast growing mutants to have a shorter C+D period that compensates for their faster growth rate. Future studies on these fast-growing mutants could be enlightening.

The relative timings of nucleoid separation and cell constriction are independent of growth rate (Fig EV4B, C). The absence of correlation between growth rate and the relative timing of these cell cycle events was also observed for the wild-type strain when the growth rate was varied by changing the composition of the growth medium (Den Blaauwen et al, 1999). Collectively, our findings show that the cell can accommodate a large range of sizes and relative timings of nucleoid segregation and cell division with no effect on growth rate, and vice versa. This flexibility may offer greater evolvability of cellular dimensions and cell cycle progression.

The complexity of cellular systems can sometimes be reduced to simple quantitative relationships, which have been very useful in identifying the governing principles by which cells integrate various processes (Scott & Hwa, 2011). Our correlation analysis identified a strong linear relationship between nucleoid size and cell size. This remarkable scaling property is independent of growth rate and holds across the wide range of cellular perturbations present in the ~4,000 deletion strains examined in this study (Fig 8C and D). This result draws a striking parallel with the 100-year-old observation that nucleus size scales with cell size in eukaryotes (Conklin, 1912), an empirical relationship that has been reported for many eukaryotic cell types since (Vukovic et al, 2016). This suggests a universal size relationship between DNA-containing organelles and the cell across taxonomic kingdoms, even for organisms that lack a nuclear envelope.

Our information-theoretic Bayesian network analysis (Fig 8) enabled us to go beyond pairwise correlations by integrating the complex set of interdependencies between cell morphogenesis, growth and cell cycle events. This analysis unveiled an unexpected connection between average cell size and the relative timings of nucleoid separation and cell constriction through nucleoid size across thousands of genetic perturbations (Fig 8E and F). This finding suggests that the size of the nucleoid is an important element of the coordination mechanism between cell morphogenesis and the cell cycle.

## Materials and methods

### Bacterial strains and growth conditions

The Keio collection contains 3,787 annotated single-gene in-frame deletion strains, 412 strains (referred to as JW strains) with kanamycin cassette inserted at unknown locations, and the remainder (28) were repeats (Baba et al, 2006).

All strains, including *E. coli* K12 BW25113 (Datsenko & Wanner, 2000) and derivatives (strains from the Keio collection), as well as *E. coli* K12 MG1655 and isogenic *ftsA\** WM1659 strains (Geissler et al, 2007) were grown in M9 medium (6 g/L Na_2_HPO_4_·7H_2_O, 3 g/L KH_2_PO_4_, 0.5 g/L NaCl, 1 g NH_4_Cl, 2 mM MgSO_4_, 1 µg/mL thiamine) with 0.2% glucose as the carbon source and supplemented with 0.1% casamino acids.

### Screening set-up and microscopy

All *E. coli* strains were grown overnight at 30°C in 96-well plates in M9 supplemented with 0.1% casamino acids, 0.2% glucose and kanamycin (30 µg/mL). Cultures were diluted 1:300 in 150 µL of fresh M9 medium supplemented with 0.1% casamino acids and 0.2% glucose, and grown in 96-well plates at 30°C with continuous shaking in a BioTek plate reader. DAPI was added to the cultures to a final concentration of 1 µg/mL 15 to 20 min prior imaging. All (parent and mutant) strains were sampled within a very narrow range of OD_600nm_ (0.2 ±0.1; min = 0.108; max = 0.350) corresponding to the exponential growth phase (Appendix Fig S1D). We did not detect any trend between morphological/cell cycle features and the OD_600nm_ at which each culture was sampled. Cells were deposited (0.5 µL per strain) on a large, 0.75-µm thick, M9-supplemented agarose pads with a multichannel pipet. The pads were made by pouring warm agarose containing supplemented M9 medium between a (10.16 × 12.7 × 0.12 cm) glass slide and a (9.53 × 11.43 cm) n° 2 cover glass (Brain Research Laboratories, Newton, MA, USA).

Microscopy was performed on an Eclipse Ti-E microscope (Nikon, Tokyo, Japan) equipped with Perfect Focus System (Nikon, Tokyo, Japan) and an Orca-R2 camera (Hamamatsu Photonics, Hamamatsu City, Japan) and a phase-contrast objective Plan Apochromat 100x/1.45 numerical aperture (Nikon, Tokyo, Japan). The initial field of view for each strain was chosen manually and 9 images were taken automatically over a 3×3 square lattice with 200 nm step, using 80 ms exposure for phase contrast and 600 ms exposure for the DAPI channel using Nikon Elements (Nikon, Tokyo, Japan).

### Image processing

Cell outlines were detected using the MicrobeTracker software (Sliusarenko et al, 2011). All data processing was then performed using MATLAB (The MathWorks Inc., Natick, MA, 2000). Custom-built codes were used to automate the aggregation of data from the cell outlines of all the strains. Data are available in Dataset EV1-2.

For cell and nucleoid detections, we consistently used the same parameters (See Appendix for parameters). In order to avoid unnecessary bias in the cell outlines, the parameters defining the initial guess for the cell contour fit were set to intermediate values, while the parameters constraining the fit of the final outline were set to negligible values. For example, we increased the *fsmooth* parameter value to 100 in order to capture both short and long cells, and we set the width spring constant parameter *wspringconst* to 0 so as to avoid biasing the cell width estimate toward the initial guess value. The edges in the DAPI fluorescence signal were detected with Oufti’s objectDetection tool (Paintdakhi et al, 2016) which is based on a Laplacian of Gaussian filtering method that takes into account the dispersion of the point spread function (PSF) of our microscopy setup at a wavelength of 460nm (input parameter *σ*_*PSF*_ set to 1.62 pixels).

### Rifampicin run-out experiments

The number of ongoing replication cycles was examined in run-out experiments (Skarstad & Katayama, 2013). BW25113 cells were grown at 30°C either in M9 glycerol or M9 medium supplemented with 0.1% casamino acids, 0.2% glucose and 1 µg/mL thiamine (as for the Keio screen described here). Cells were grown up to exponential phase and then treated for 3 h with 30 µg/mL cephalexin and 300 µg/mL of rifampicin prior to overnight fixation in 70% ethanol at 4 °C. Cells were washed twice with phosphate buffered saline and then stained with DAPI (1µg/mL) prior to imaging on a PBS-containing agarose pad. In M9 glycerol medium, *E. coli* BW25113 cells do not start a new round of replication before the previous one ended (Cooper & Helmstetter, 1968; Wang et al, 2011). This growth condition was used as a control to estimate the DAPI intensity corresponding to 1 and 2 genome equivalents.

## Data analysis

### Dataset curation – Support Vector Machine (SVM) model

Due to the size of the dataset (>1,500,000 cells detected globally), we adopted an automated approach to identify poorly (or wrongly) detected cells across the entire dataset. We developed an SVM model based on 16 normalized features: cell length, cell width, cell area, cell volume, cell perimeter, cell constriction degree, division ratio, integrated phase signal, integrated DAPI fluorescence signal, mean cell contour intensity in phase contrast, variability of cell width along the cell, nucleoid area, single cell nucleoid variability, cell circularity (2\**π*\*cell area/(cell perimeter)^2^), nucleoid intensity and number of nucleoids. We trained a binary classifier (positive or negative) over wild-type strain replicates as well as 419 mutants with the most severe morphological defects prior to data curation. We visually scored 145,911 cells and used 30% of them (43,774) to train the model. The model was evaluated using a k-fold cross-validation approach, leading to a generalized misclassification rate of 10%. We used the remaining 70% of the data set (102,137 cells) to validate the model. This SVM classifier achieves a balanced classification rate of 84% and features an AUROC of 0.94 (Appendix Fig S1E). Furthermore, the resulting group of false negatives was not significantly different from the true positives (Appendix Fig S1F and G), indicating that the classification did not introduce a bias by excluding a specific class of ‘good’ cells from the analysis.

### Data processing

For each feature, we checked and corrected for any bias associated with plate-to-plate variability, differences in position on the 96-well plates, timing of imaging and optical density of the culture (Appendix Fig S2-S4). For each plate, we set the median values of each feature, *F*, to the median feature value of the parental strain.

The *F* values were transformed into normalized scores by a transformation akin to a z-score transformation but more robust to outliers.
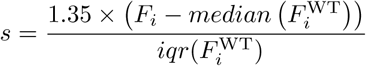

where *F*_*i*_ is the corrected value for the mutant strains for feature *i*, 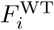 is the value for the wild-type strain for feature *i*, and iqr stands for interquartile range. As the interquartile range of normally distributed data is equal to 1.35 times their standard deviation, we scaled the score by this factor so as to express the scores in terms of standard deviations away from the median.

The temporal biases for the fraction of cells committed (or not) to division and the fractions of cells with 1, 2 or more nucleoids were corrected using a Dirichlet regression to maintain the relative proportions between classes (Appendix Fig S3) (Maier, 2014). In an exponentially expanding population of growing *E. coli* cells at steady state, the fraction of cells in the population before the occurrence of a cell cycle event is related to the cell age at which this event occurs by a monotonic relationship (Collins & Richmond, 1962). The proportions of cells at different cell cycle stages are therefore used to infer the relative proportions of different cell cycle stages and the cell age at which a specific cell cycle event occurs (Collins & Richmond, 1962; Powell, 1956). These inferences rely on the assumption of ergodicity, which is assumed in the analysis of population-based experiments such as cell cycle studies by flow cytometry or marker frequency analysis (Grangeon et al, 2015; Kafri et al, 2013). One limitation of this approach is that only relative timings or proportions can be inferred, and no conclusions should be drawn on the absolute duration of the different periods without any other hypothesis. For instance, a reduction of the fraction of cells with one nucleoid could result either from an actual reduction in the cell cycle period associated with one nucleoid, or from a lengthening of the period associated with cells with two or more nucleoids. In both cases, we can, however, conclude that the separation of nucleoids happens earlier in relative cell cycle units.

### Data exploration, dimensionality reduction and clustering

A similarity measure between strains was needed to identify and separate different phenoprints. Pearson correlations or Euclidean distances classically provide such similarity measures, and Principle Component Analysis (PCA) and/or hierarchical or k-means clustering are often used. However, PCA tends to explode datasets and Pearson correlations do not always reflect the desired type of similarity. As an extreme example, consider two strains with two phenoprints that are proportional, one with values within a very small score range, such as [-1 1], while the other with score values spanning the [-10 10] range. These two strains will get a maximal similarity measure through a correlation analysis, despite the fact that the first strain is wild-type-like while the other is an outlier. Instead we chose to use a recently described algorithm, called t-distributed Stochastic Neighbor Embedding, or t-SNE (van der Maaten & Hinton, 2008), to project our multidimensional datasets in 2 dimensions and generate, at the same time, similarity measures between strains. t-SNE estimates low-dimensional space distances between points based on their similarity, as opposed to dissimilarity as in the case of PCA, thereby highlighting local similarities rather than global disparities.

We used the stochastic nature of the t-SNE algorithm to evaluate the robustness of the resulting projection by repeating the procedure multiple times (*n* = 100 for each tSNE map). We coupled this dimensional reduction procedure with a density-based clustering algorithm, dbscan (Ester et al, 1996), to group strains with similar phenoprints. The two input parameters of the dbscan algorithm, *E* and minPoints, were optimized so as to generate a maximum number of islands without separating the bulk of WT strains in two or more islands. Islands include strains that clustered together more than 90% of the time.

The convergence of the dimensionality reduction was verified by repeating the tSNE dimensionality reduction on sub-samples of the initial dataset (1,225 × 21 matrix). In this approach akin to cross-validation, we generated 50 partitions of either 1,200 or 1,201 strains, holding out disjoint sets of points, with the cvpartition built-in function in MATLAB, and repeated the dimensional reduction with the tSNE algorithm 10 times for each partition. We first compared the pairwise distances between the points in each of these 500 tSNE maps with the pairwise distances between the corresponding points in the tSNE map presented in Fig4A. The distribution of Pearson correlation coefficients between these sets of distances, calculated with a kernel density estimation function (Botev et al, 2010), is illustrated in Appendix Fig S8A (red curve). This distribution is highly similar to the distribution obtained from the repetition of the tSNE dimensional reduction on the full dataset (1,225 strains – blue curve in Appendix S8A), and suggests that the algorithm converges toward a global minimum. The bimodality in the distribution of the Pearson correlation coefficients is not due to specific sub-samples (Appendix Fig S8B) and rather reflects the stochasticity of the tSNE algorithm and the low weight carried by large distances in the map. For example, the displacement of a well isolated island relative to others may not impact strongly the minimized score of this tSNE map, but would definitely reduce the Pearson correlation coefficient between this map and the reference tSNE map presented in Fig 4A. Using the same parameters as for the full dataset (*E* = 4.9, minPoints = 3), we verified that the dbscan algorithm results in a similar clustering as in the global map. The resulting clusters for each tSNE map associated with a sub-sample of the dataset was compared with the clustering output presented in Fig 4A and B. For each reference cluster (islands in Fig 4A), we identified all the representative points of this cluster in the sub-sample tSNE map and calculated the ratio between the maximal number of these points clustering together in this sub-sample map divided by the total number of representative points of the reference cluster in this sub-sampling. This ratio is akin to the Jaccard index for each reference cluster, and all indexes for a given map were averaged to provide a score to each tSNE map. The distribution of these scores (10 scores per sub-sample corresponding to the 10 tSNE maps calculated for each sub-sample) are represented in the boxplot in Fig S8C. The average indexes are typically above 0.9, which reflects the threshold used to generate the clusters as points clustering in more than 90% of the tSNE maps. The same reproducibility of clustering in the sub-sample tSNE maps is also a good indication that the tSNE dimensional reduction converges toward an optimal embedding.

### Map exploration

Each t-SNE map is a similarity map, and can therefore be treated as a network where the nodes represent strains and the edges the Euclidean distance between strains in the tSNE map. Building up on recent quantitative network analysis tools (Baryshnikova, 2016), we calculated the local enrichment in the maps of different strain-associated attributes, such as COG and GO terms. Briefly, the sum of the attributes in a local area (within a radius around each point, defined as the 1-percentile of the distribution of all the pairwise distances between points) was compared to a background score (defined as the average score obtained over 1000 identical maps with randomly permutated attributes) with a hypergeometric test. The significant local enrichments were considered at a threshold of 0.05 after adjusting for false discovery rate correction with the Benjamini-Yekutieli (BY) procedure, taking into account dependencies between tests (Benjamini & Yekutieli, 2001). The SAFE algorithm proposed by Baryshnikova (Baryshnikova, 2016) was implemented as a MATLAB function mapEnrich.m (see computer code EV1A).

### Cluster of orthologous gene enrichment analysis

We associated *E. coli* BW25113 genes with COGs using the web server (Van Domselaar et al, 2005). The enrichment analyses were performed using a custom-built algorithm in MATLAB based on a two-tailed hypergeometric test to compute p-values, which were subsequently adjusted with the Benjamini-Hochberg (BH) False Discovery Rate procedure (Benjamini & Hochberg, 1995) (Computer code EV1B). Because the COG categories are largely independent, we did not consider any correction for the dependence between tests.

### Gene ontology analysis

We used ontologies from the Gene Ontology website (http://www.geneontology.org/ontology/gene_ontology.obo, version 2016-05-27) (Ashburner et al, 2000), and annotations were obtained from EcoCyc for *E. coli* strain MG1655 (Keseler et al, 2013). Analysis was performed using a MATLAB custom-built algorithm that includes a hypergeometric test to compute p-values that were subsequently adjusted with the BY False Discovery Rate procedure (Benjamini & Yekutieli, 2001) (Computer code EV1B).

### Bayesian network

The Bayesian network presented in Fig 8 was generated in R with the bnlearn package (Scutari, 2010), using the ARACNE algorithm as described in (Margolin et al, 2006). The network was bootstrapped 200 times, and all the edges were identified in more than 70% of the networks. We assessed the strength and the origin of collinearity among features using Belsley diagnostic method (Belsley et al, 1980), with the built-in collintest.m function in MATLAB. We excluded features associated with a ‘condition number’ above the classical threshold of 30.

### Calculation of confidence intervals

As the true correlation between two normally distributed variables approaches 1 or −1, the probability density distribution of the estimated correlation becomes highly skewed and far from normal. For Pearson correlations (*ρ*), this distribution can be transformed using Fisher z-transform to approximate a normal distribution for any true correlation value *r* (Fisher, 1915):
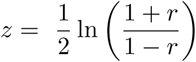

The variable *z* follows a normal distribution with mean 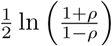 and standard error 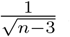, where *ρ* and *n* are the estimated Pearson correlation value and the sample size, respectively.

Using this transformation, the limits of the distribution covering the 95% most probable values of the distribution can be calculated using the 2.5 and 97.5 percentiles of the normal distribution centered on 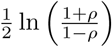 with standard deviation 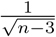.

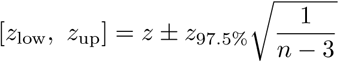

The inverse transformation on these confidence boundaries on z is then used to calculate the 95% confidence upper and lower limits for the estimated correlation value *ρ*.

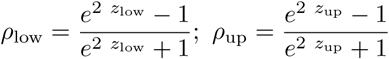

Unfortunately, a similar approach cannot be employed for an estimated Kendall correlation (*τ*) value (Long & Cliff, 1997). To generate 95 % CIs, we used a bootstrapping approach (Efron & Tibshirani, 1993). Correlation values were calculated between 5000 resampled (with replacement) variable pairs. The 2.5 and 97.5 percentiles of the obtained distribution of correlation values were subsequently taken as the respective lower and upper boundaries of the bootstrapped 95 % CIs.

## Data representation

All graphs were generated using MATLAB, except for the networks in Fig 4C and Fig 8A panels, which were created using Cytoscape v3.2 (Shannon et al, 2003) and the Rgraphviz package in R (Hansen et al), respectively. For Fig 4C, we used the built-in edge-weighted, spring embedded algorithm in Cytoscape. We considered the pairwise Euclidean distances between the 9 strains of island 22 as the weights of the edges connecting the nodes (or strains).

The density scales in scatter plots represent the number of points around each point in a radius equal to the 0.03 percentile of the pairwise distances distribution.

The WT isocontours representing the 0.5, 0.75 and 0.95 probability envelopes for the 240 WT replicates were calculated using a 2D kernel density estimation function over a 128-by-128 lattice covering the entire set of points. The bandwidth of the kernel was internally determined (Botev et al, 2010).

## Simulations of cell length distributions

Cell length distributions at any given cell age were assumed to be log-normally distributed with different dispersion values. The CV of the distribution for the WT strain (CV = 0.11) was previously experimentally determined (Campos et al, 2014). The cell length distributions at 100 different ages equidistantly distributed between 0 (birth) and 1 (division) were convolved with the cell age distribution, assuming an exponentially growing culture, *P r*(*age*) = 2^*−age*^.

## Acknowledgements

We are grateful to the Yale E. coli Genetic Stock Center for providing a large number of strains. We also thank Pr. William Margolin for the kind gift of the *E. coli* MG1655 strain and the *ftsA\** derivative. This work was partly supported by the National Institutes of Health (R01 GM065835 to C.J.-W.). We also thank the Jacobs-Wagner laboratory for fruitful discussions and for critical reading of the manuscript. M.C. was partly funded by a fellowship from the “Fondation pour la Recherche Médicale” (ARF20160936199). C.J.-W. is an investigator of the Howard Hughes Medical Institute.

## Author contributions

C.J.-W, and M.C. designed experiments. G.S.D, S.G., I.I. and M.C performed experiments. M.C. performed high-throughput imaging and statistical analyses. C.J.-W. supervised the project. C.J.-W. M.C., I.I. and S.G. wrote the manuscript.

## Dataset EV1. Raw data

Excel file (*.xlsx) of the raw data of SVM-curated dataset prior to any correction and normalization.

## Dataset EV2. Score table

Excel file compiling all quantitative information about each strain imaged in this study, including the scores and measured values (after correction and normalization) for each morphological, growth and cell cycle feature, as well as their association to a morpho or cell-cycle island when appropriate.

## Computer Code EV1A

MATLAB-based function used to calculate local enrichments for GO terms (related to Fig 4E and Fig 5D). For each point in the tSNE map, a neighborhood was defined by taking all the points closer than a distance *r* defined as the 1-percentile of the distribution of pairwise distances among all points in the tSNE map. For each GO term, a hypergeometric test was performed within each neighborhood. The resulting p-values are adjusted to control the false discovery rate using the BY procedure (Benjamini & Yekutieli, 2001).

## Computer Code EV1B

MATLAB-based function dedicated to false discovery rate control. The function takes a set of p-values, a threshold and a procedure as inputs, and provides a set of adjusted p-values (q-values) and a logical array indicating which q-value is below the threshold used as an input. The two possible procedures to control the FDR correspond to the algorithms known as the BY and BH procedures, taking into account dependencies between tests (BY procedure) (Benjamini & Yekutieli, 2001), or not (BH procedure) (Benjamini & Hochberg, 1995). The p-values associated with COGs enrichments were corrected with the BH algorithm, which is not taking into account dependencies between tests because COGs are vastly non-overlapping. The p-values associated with GO term enrichments were corrected with the BY algorithm because GO terms can encompass overlapping sets of genes.

### Expanded View Figures

**Figure EV1.**
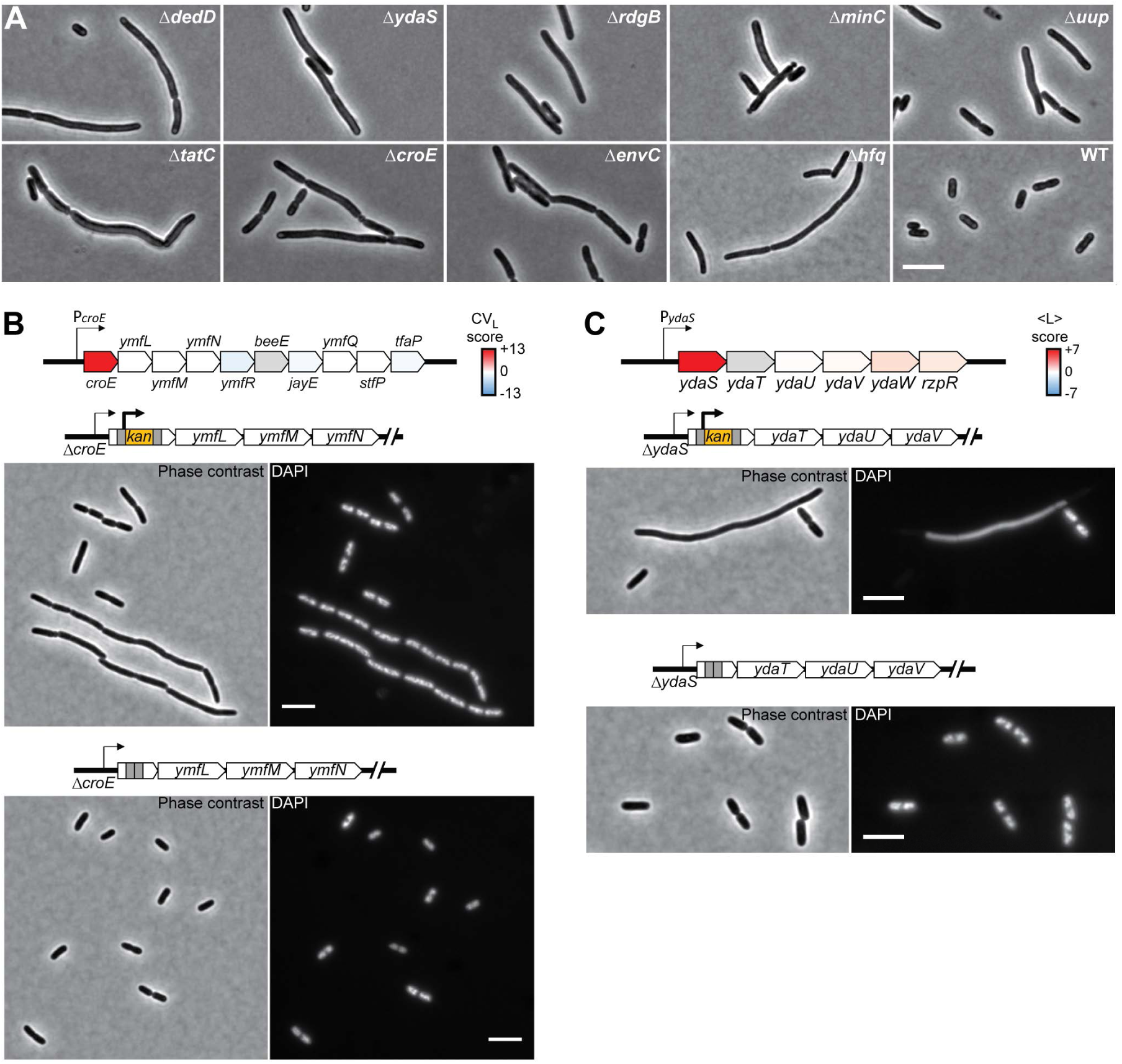
Filamentous mutants. **A.** Representative phase-contrast images of the mutants forming the island 22, together with the parental strain BW25113 (WT) for comparison. **B.** Effect of the kanamycin-resistance cassette on the phenotype of the ∆*croE* strain. The schematic at the top shows the color-mapped score of CV of cell length for the deletion of each gene of the *croE* operon. Below are phase-contrast and fluorescent images of DAPI-stained cells of the ∆*croE* strain carrying the kanamycin resistance cassette (top) or after the removal of the cassette (bottom). The *ymfN* locus has been re-annotated as two separate genes (*oweE* and *aaaE*), and the Keio deletion strain of *ymfN* carries the deletion of these two contiguous genes. **C.** Effect of the kanamycin-resistance cassette on the phenotype of the ∆*ydaS* strain. The schematic at the top shows the color-mapped score of the mean cell length for each gene of the *ydaS* operon. Below are phase-contrast and fluorescent images of DAPI-stained cells of the ∆*ydaS* strain carrying the kanamycin resistance cassette (top) or after the removal of the cassette (bottom).

**Figure EV2.**
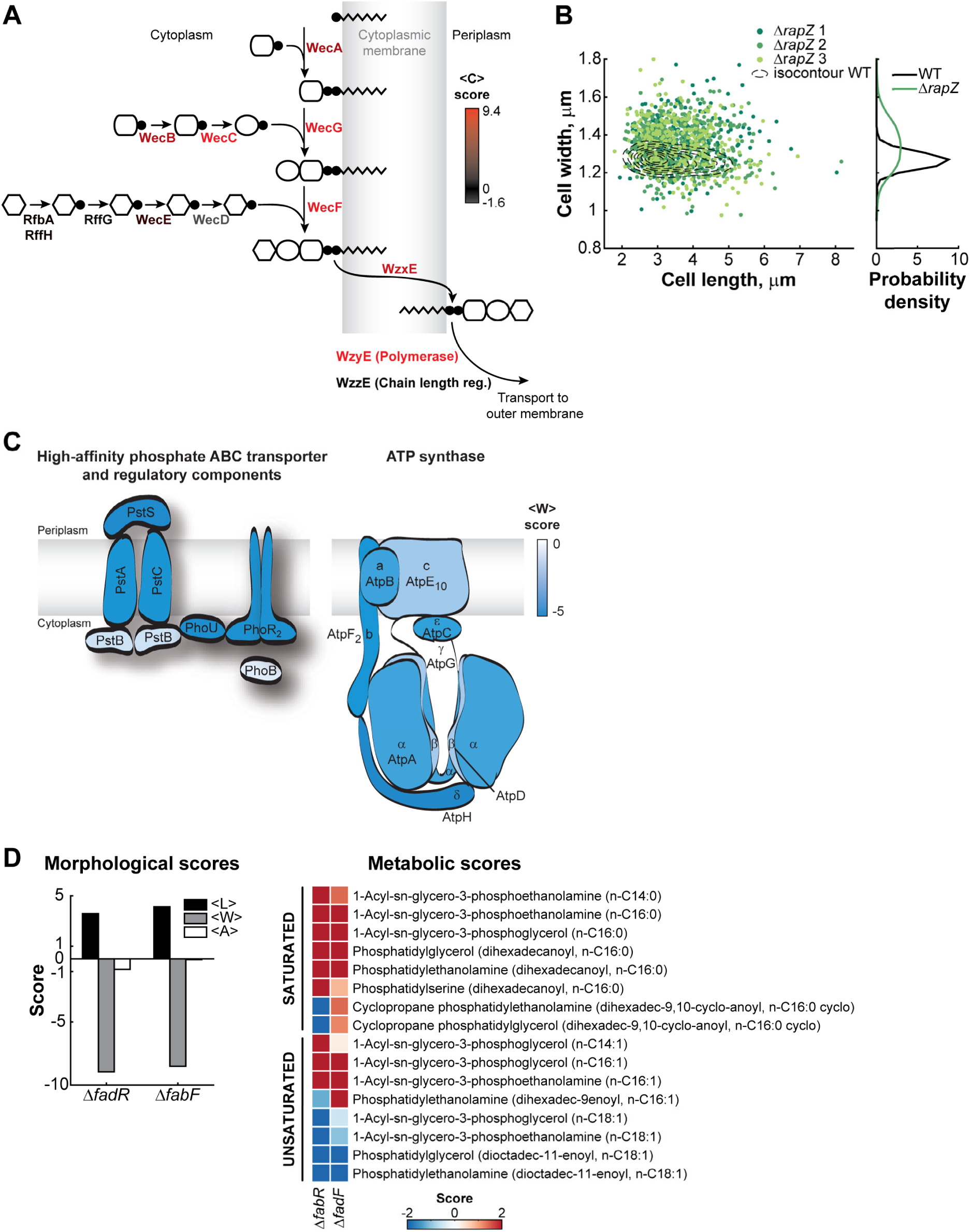
Specific pathways associated with impaired cell morphology. **A.** Schematic of the ECA biosynthetic pathway in which each gene name has been colored by the severity of the mean circularity (<C>) phenotype. **B.** Scatter plot of cell width versus cell length for three independent liquid cultures of the ∆*rapZ* strain (*n* = 564, 268 and 343 cells) in well-agitated test tubes. The dotted lines represent iso-contours of a 2D histogram of cell length and cell width for the parental strain (WT, *n* = 1,045 cells). The cell width distributions of the WT and ∆*rapZ* strains are represented on the right of the scatter plot (all three replicates for the ∆*rapZ* strain were pooled together). **C.** Schematics of the ATP synthase, the high-affinity ABC phosphate transporter and its related proteins. Proteins and subunits have been colored according to the severity of the mean cell width (<W>) phenotype in the corresponding gene deletion strains. **D.** Bar graph showing the mean cell length (<L>), width (<W>) and area (<A>) scores for the ∆*fadR* and ∆*fabF* deletion strains. The scores for saturated and unsaturated phospholipid metabolites detected in these strains by Fuhrer et al. (2017) are represented as a heatmap.

**Figure EV3.**
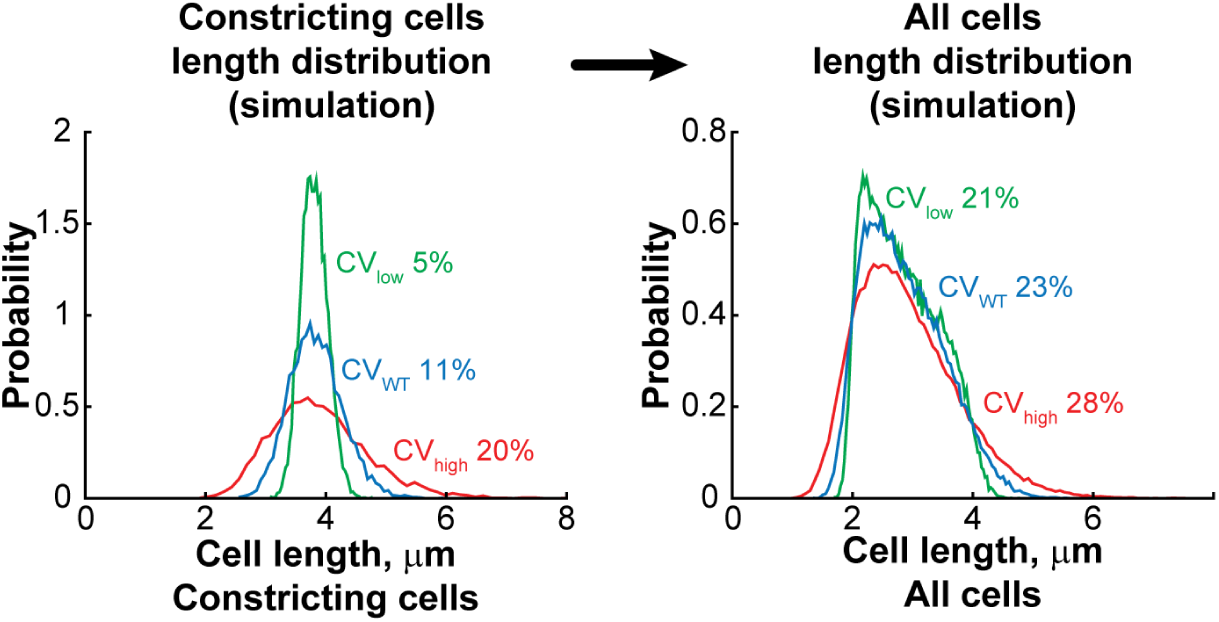
Simulation showing that the cell length variability of the entire population can mask abnormal cell length variability at a specific cell cycle period. Cell length distributions were simulated over different ranges of cell ages (see Materials and methods). The cell length distribution of constricting cells was determined by summing the cell length distributions of all cells of age > 0.8, assuming different CV of the cell length distribution (0.05, 0.11 and 0.2) at a specific age. The cell length distribution of the whole population was determined by summing the distributions at all ages, from birth to division.

**Figure EV4.**
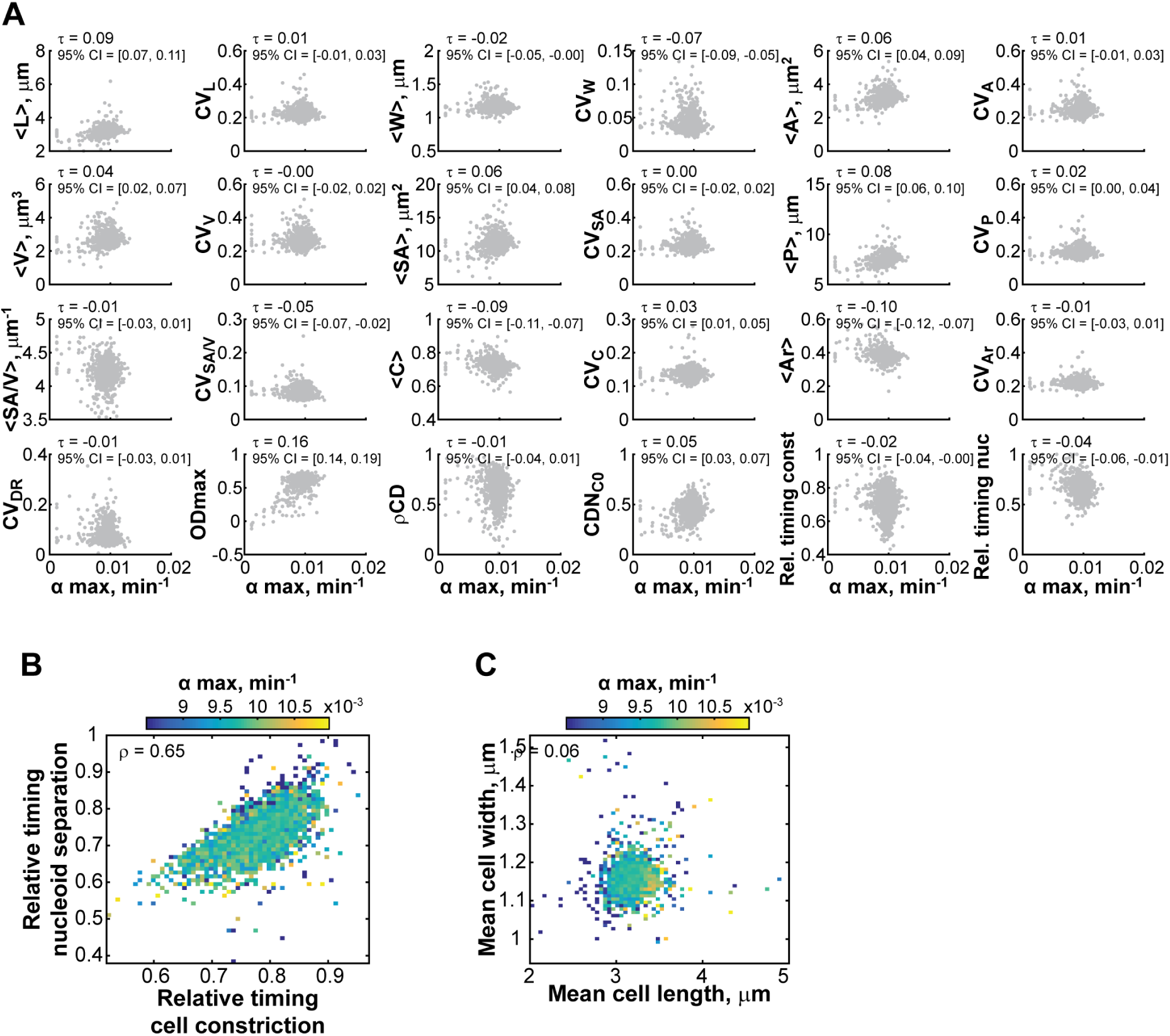
Growth rate correlates poorly with morphological and cell cycle features. **A.** Scatter plots of morphological and cell cycle features versus *α*_max_. Each grey dot represents one Keio strain. The Kendall correlation coefficient *τ* and the associated 95% confidence interval are reported for each pair of features. The confidence intervals were calculated by bootstrapping the correlation 5000 times and taking the 2.5 and 97.5 percentiles of the resulting distribution. The Kendall ranked correlation *τ* was selected over Pearson correlation because of the heavily asymmetric left tail in the distribution of *α*_max_. **B.** Heatmap showing the mean growth rate value for data binned by the relative timings of cell constriction and nucleoid separation. **C.** Same as **B**, except for data binned by mean cell length and mean cell width.

**Figure EV5.**
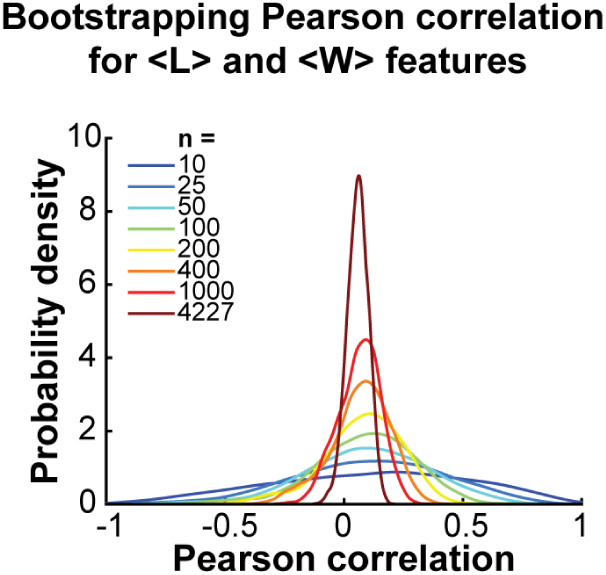
The sample size affects the confidence in the estimation of the correlation coefficient. The scores for the <L> and <W> features were sampled 5,000 times independently with replacement with different sample sizes (n = 10, 25, 50, 100, 200, 400, 1,000 or 4,227), and the Pearson correlation coefficient was calculated at each sampling. The distribution of the 5,000 correlation values obtained for each sample size is represented with a different color. The probability density distributions were estimated using a kernel density estimation method (Botev et al, 2010). The plot shows that a small sampling size, such as n =10, results in a wide distribution of possible Pearson correlations. Increasing the sample size narrows down the distribution, increasing the confidence in the obtained correlation value.

### Appendix Figures

**Appendix Figure S1.**
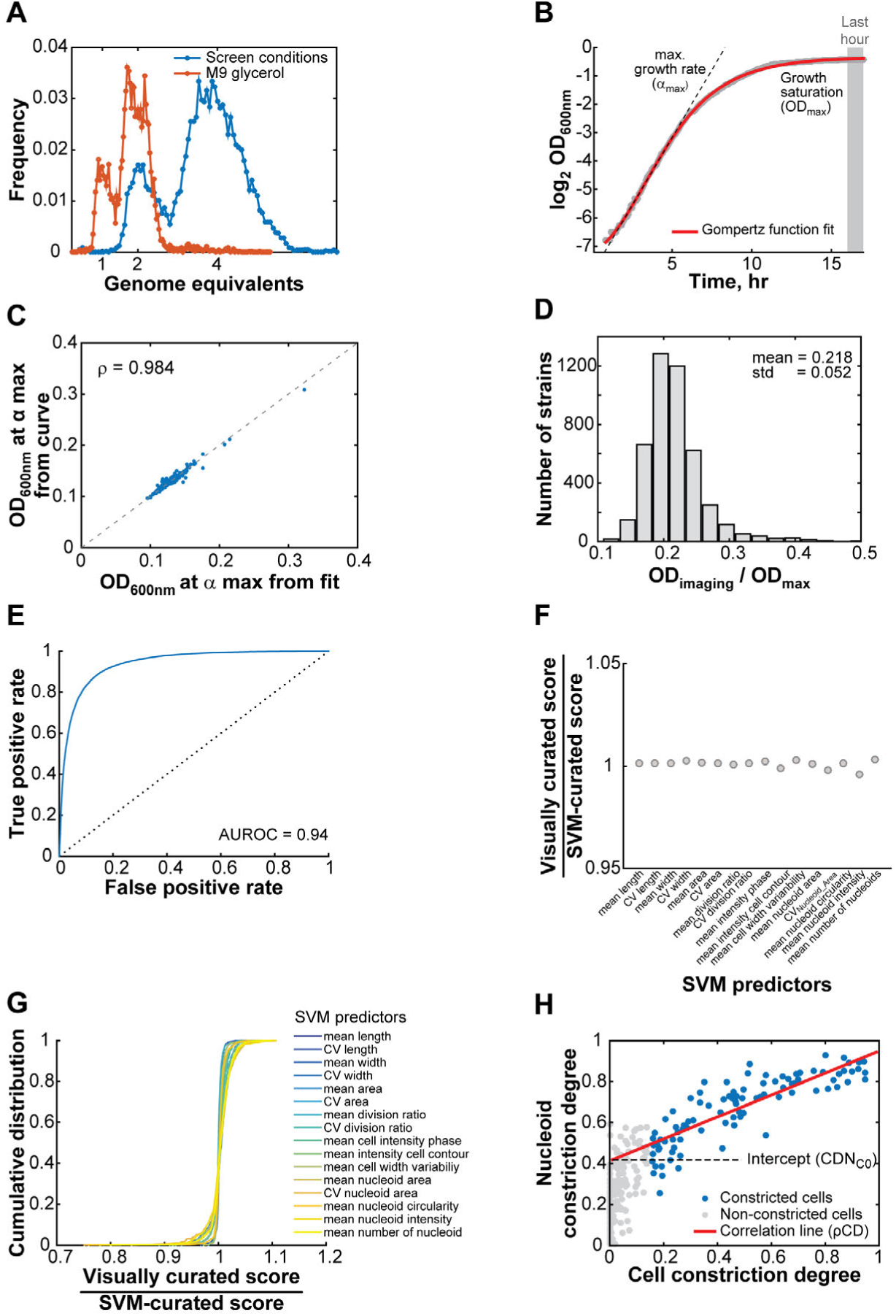
Feature determination and SVM model validation. **A.** Distribution of genome equivalents per cell determined by the signal intensity of DAPI DNA stain after a replication run-out experiment. BW25113 cells were grown at 30^*◦*^C either in M9 glycerol (poor nutrient condition – orange curve) or under the same growth conditions as for the screen (M9 glucose supplemented with casamino acids, a richer nutrient condition – blue curve), and then treated with rifampicin and cephalexin for 3 h to block cell division and prevent new rounds of DNA replication. Cells grown in M9 glycerol undergo a single cycle of DNA replication per division cycle (Cooper & Helmstetter, 1968; Wang et al, 2011). After rifampicin and cephalexin treatment, these cells contained either one or two chromosomes, and displayed a fluorescence level with DAPI corresponding to 1 and 2 genome equivalents. This calibration was used to determine the number of genomes per cell for cells growing under the richer nutrient conditions used for the screen. **B.** Typical growth curve represented as the log_2_ of OD_600nm_ as a function of time. The red line shows the best Gompertz fit to the curve (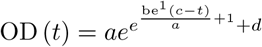 where *t* is time and *a*, *b*, *c* and *d* are the four fitted parameters). The dotted line highlights the segment of maximal growth. The OD_600nm_ was averaged over the last hour of growth (gray box) to estimate the saturation level of the culture (OD_max_). **C.** Scatter plot showing the relationship between the OD_600nm_ at *α*_max_ calculated from the growth curve and fitted curve. The correlation between both optical densities is high (*ρ* = 0.984, 95% CI [0.983, 0.985]). **D.** Histogram showing the ratio between the optical density at the time of sampling (OD_imaging_) and the optical density at saturation of the culture (OD_max_) measured from the growth curve. The mean value of this ratio is low (0.218 ± 0.052), indicating that Keio strains were imaged early in their population growth cycle. **E.** AUROC curve (performance curve) related to the SVM model on the dataset that was not used to train the model. The dotted line y=x represents the expectation from a random classification. **F.** Ratios between the mean values of each predictor for the visually-curated and SVM-curated datasets, showing the lack of bias. **G.** Cumulative distributions of mean score ratios between visually- and SVM-curated datasets of mean and CV predictor values for the 419 strains with the most extreme phenotypes. The steepness of the curves shows that the SVM model performed well, even for strains with strong phenotypic defects. **H.** Plot showing how two cell cycle features, the correlation between the degrees of constriction of the nucleoid and of the cell (*ρ* CD) and the projected degree of nucleoid constriction at the onset of cell constriction (CDN_C0_), were calculated using a WT culture as an example. The degree of constriction for both the nucleoid and the cell (considering only cells with a degree of cell constriction over 15%) were used to calculate their Pearson correlation coefficient (*ρ* CD). The correlation coefficient can be interpreted as the slope of the line passing through the data, and the intercept of this line with the y-axis provides the average degree of constriction of the nucleoid at the onset of cell constriction (CDN_C0_).

**Appendix Figure S2.**
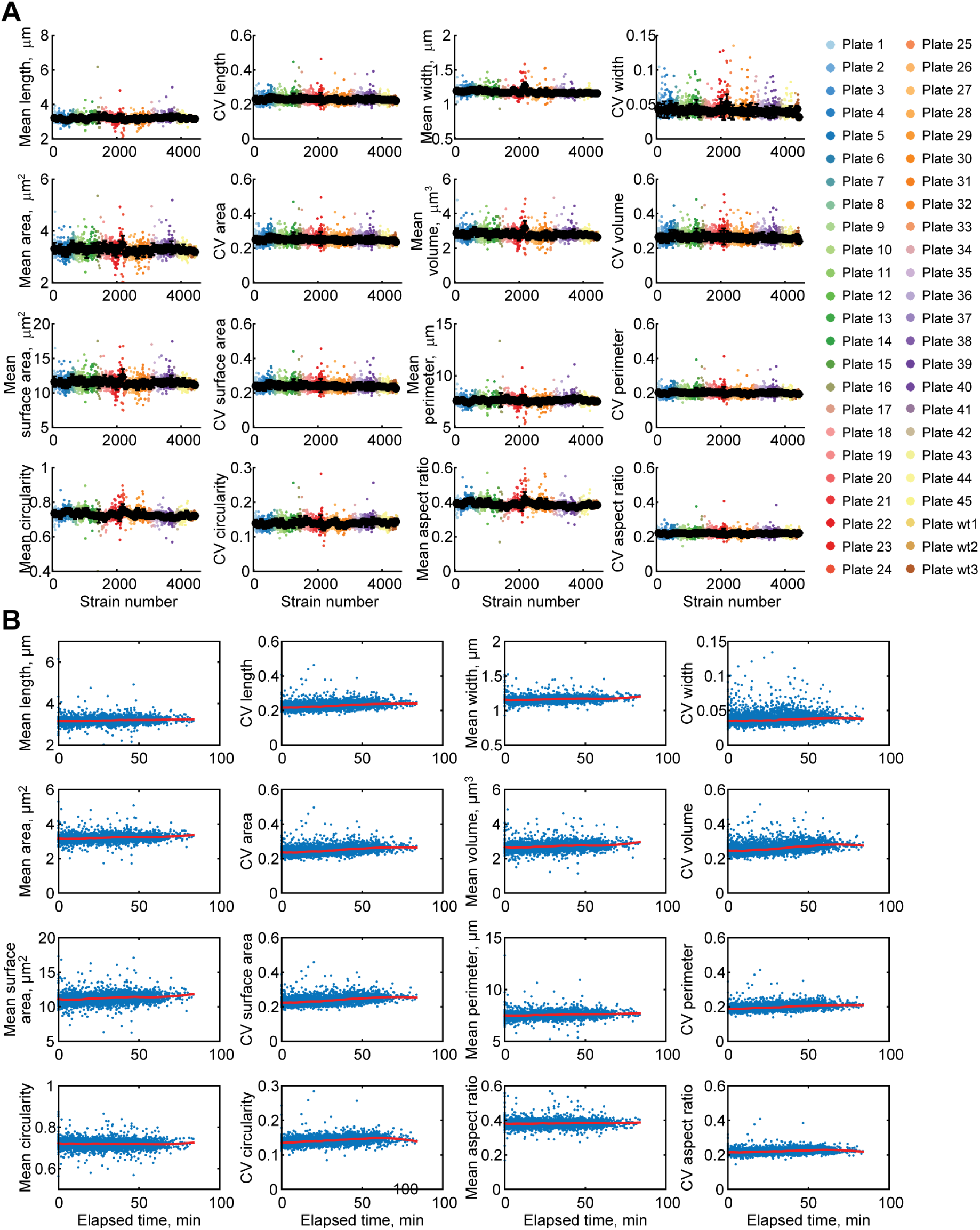
Evaluation of positional and temporal biases related to imaging. **A.** Plate-by-plate normalization. Each plate is color-coded according to the 96-well plate number. The black dots represent the mean feature value per plate, ± standard deviation. **B.** Scatter plots for each feature of all strains as a function of the time elapsed since spotting cells on the pad for imaging. The red line in each graph represents the smoothing spline (calculated with a span of 20%) that was used to correct any temporal bias.

**Appendix Figure S3.**
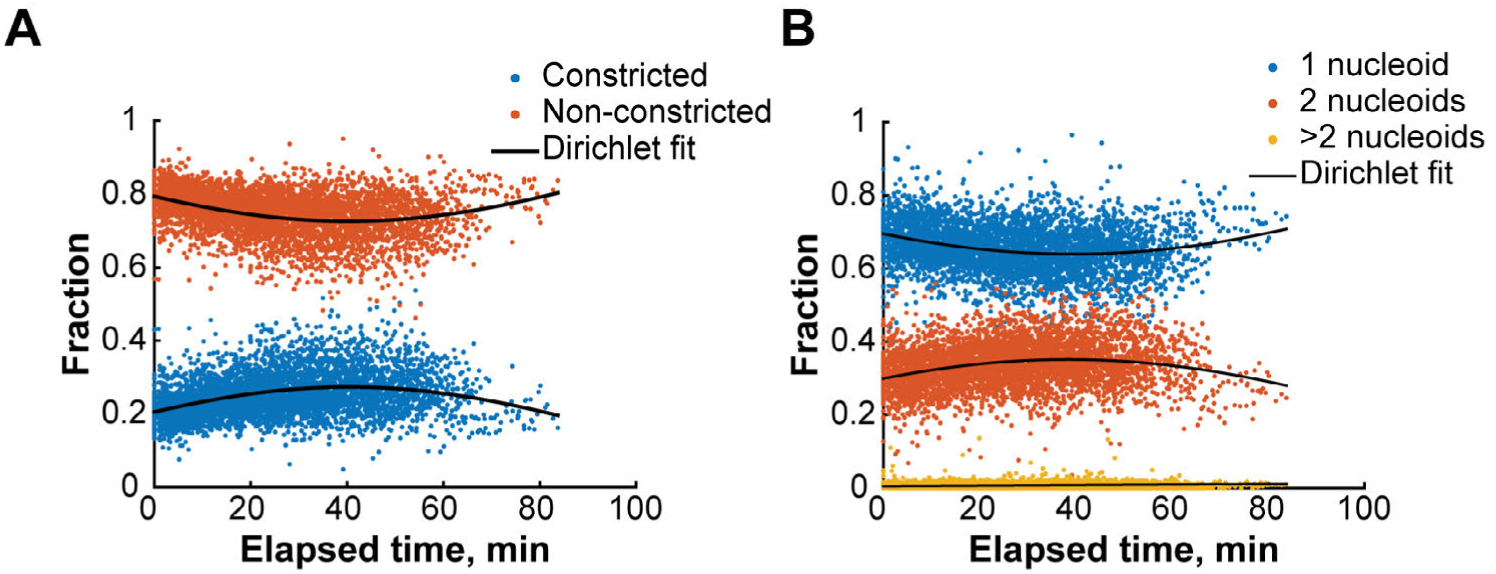
Temporal bias correction for proportional features. **A.** Scatter plot showing the proportions of constricting (blue) and non-constricting cells (red) for all strains (n = 4,227) as a function of the time elapsed between the time the cells were spotted on the pad and the time they were imaged. The solid black lines represent the correction factors over time, derived from a quadratic form Dirichlet regression to the data. The Dirichlet regression allows for the maintenance of the additivity of the proportion (Maier, 2014). **B.** Same as in **A** for the complementary proportions of cells with 1 (blue), 2 (red) or more than 2 (yellow) nucleoids. The fitted model was quadratic for the first two features (1 and 2 nucleoids) and linear for the cells with more than 2 nucleoids.

**Appendix Figure S4.**
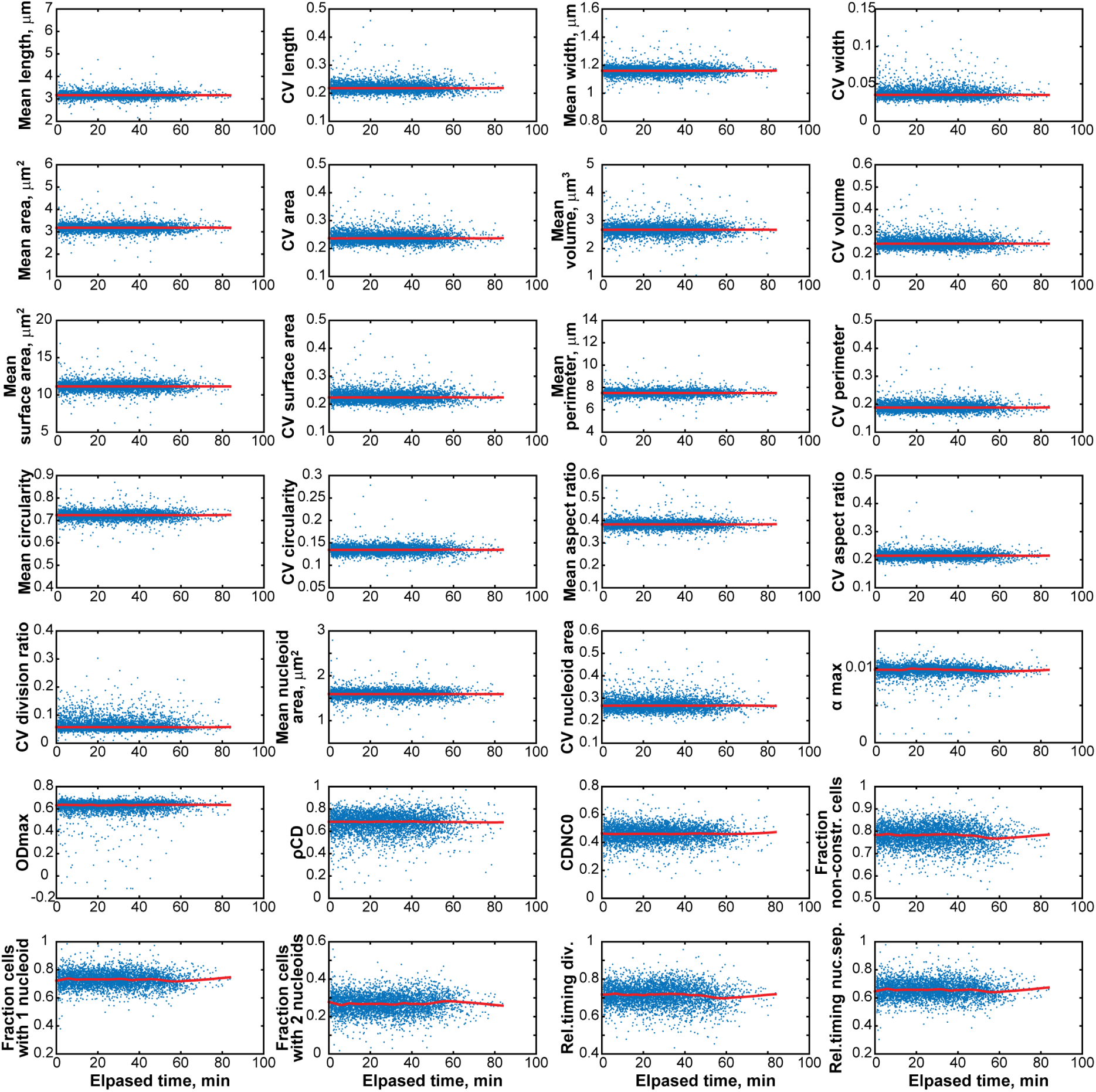
Temporal detrending. Scatter plot of the detrended data for all features. The red line in each graph represents the smoothing spline (calculated with a span of 20%). The flat profile for each of these average profiles illustrates the absence of temporal trend in the normalized data.

**Appendix Figure S5.**
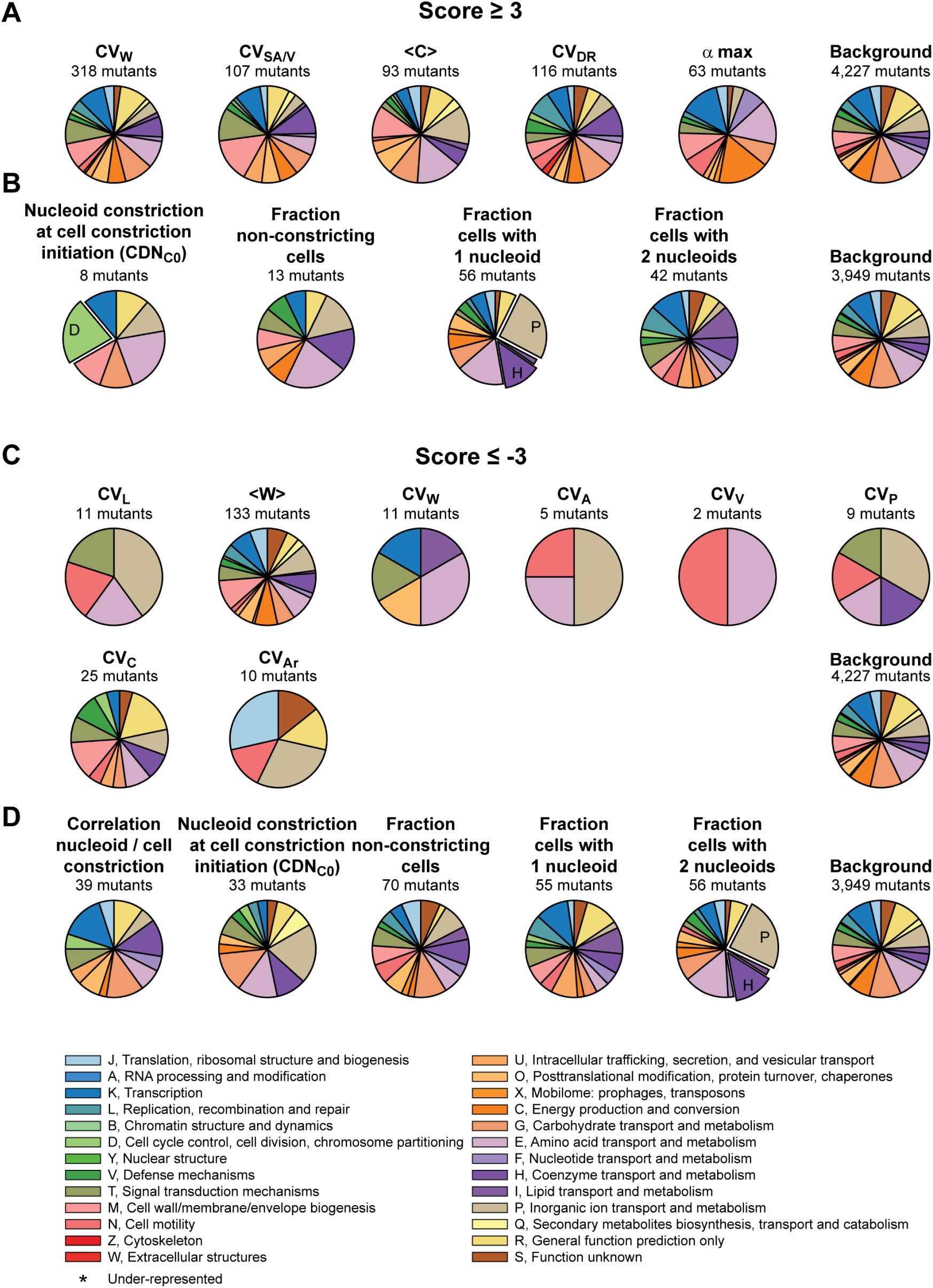
Feature-based COG distribution analysis. Pie charts representing, on a feature-by-feature basis, the relative distribution of COG categories among the gene deletion strains associated with a severe phenotype: **A.** *s* ≥ 3, **B.** *s* ≤ −3. All the features that were not included in Figure 3 are represented. The enriched COG categories are highlighted with an exploded pie sector. Enrichments with an associated (FDR corrected) q-value < 0.05 were considered significant.

**Appendix Figure S6.**
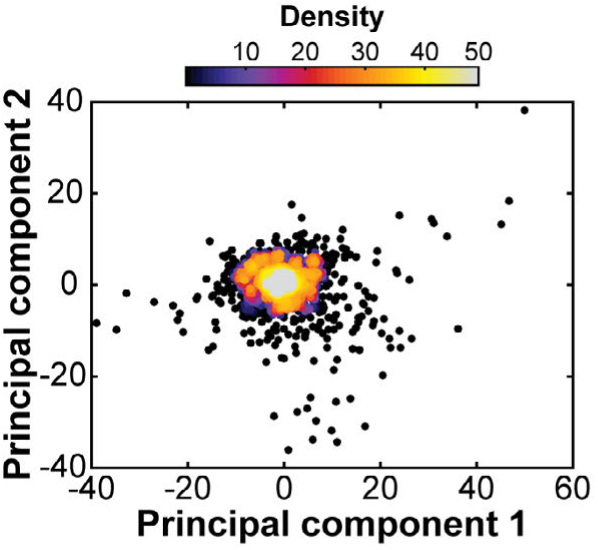
Principal component analysis of the phenoprints associated with the morphological and growth mutants. Scatter plot showing the coordinates of each strain in the first two principal components space. The density of points in any given area of the 2D space is illustrated by a color scale.

**Appendix Figure S7.**
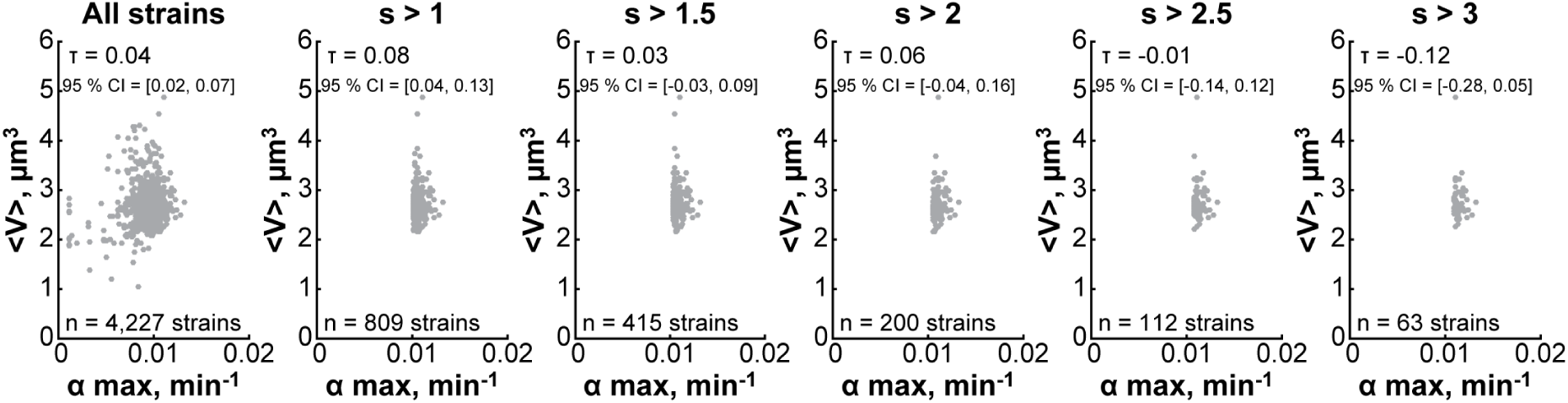
Growth rate is not predictive of cell size even for fast-growing mutants. Bootstrapped Kendall correlation values (5,000 samplings) between growth rate (*α*_max_) and mean cell volume (<V>) for all Keio strains (n = 4,227 strains) or for faster growing strains with *α*_max_ score > 1 (n = 809 strains), *α*_max_ score > 1.5 (n = 415 strains), *α*_max_ score > 2 (n = 200 strains), *α*_max_ score > 2.5 (n = 112 strains), *α*_max_ score > 3 (n = 63 strains).

**Appendix Figure S8.**
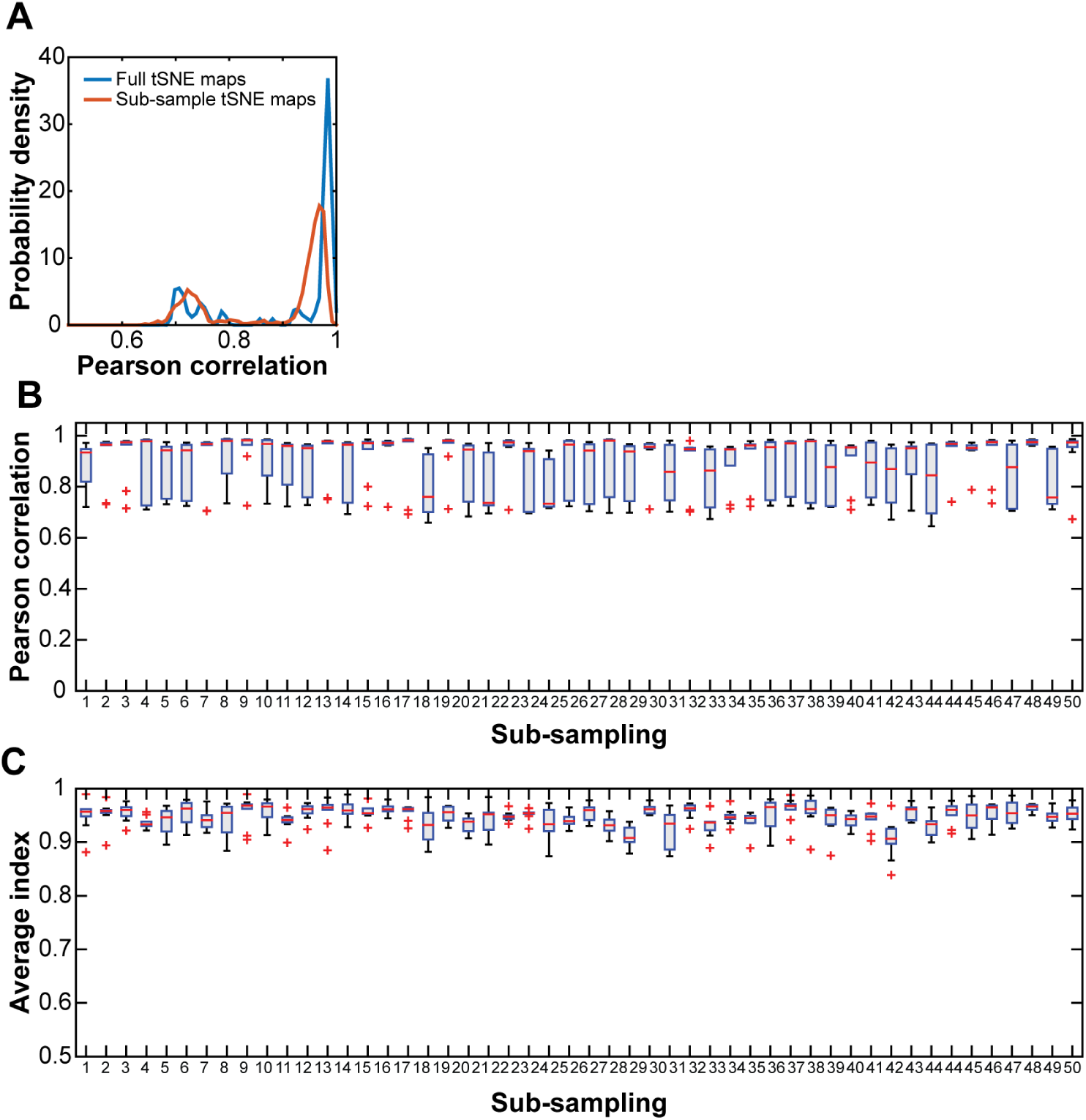
Convergence of the dimensional reduction with tSNE. **A.** Probability density distribution of the Pearson correlation coefficients between pairwise distances of the points defined in the tSNE map represented in Fig 4A and those defined in 99 other tSNE maps for the 1,225 strains that exhibit a |*s*| > 3 for at least one morphological or growth feature (blue curve). We subsampled the original dataset in 50 disjoint sets of phenoprints, and generated 10 independent tSNE maps for each one. We then calculated the Pearson correlation coefficient between the tSNE coordinates of these points in each of these 500 tSNE maps with their corresponding coordinates in the tSNE map presented in Fig 4A. The orange curve represents the probability density distribution of these correlation values. The results show that all 500 tSNE maps are very similar despite their stochastic nature, indicating convergence. **B.** Box plot showing the tight distributions of Pearson correlation coefficients between the pairwise distances between points for 10 independent tSNE maps calculated for each of the 50 sub-samples (500 tSNE maps) and their corresponding points in the tSNE map presented in Fig 4A. The blue boxes filled in gray represent the interquartile range, the red horizontal lines correspond to the median values, the whiskers cover the 2.7 standard deviation around the mean under a normality assumption, and the red crosses show the outliers beyond the limit of the whiskers. **C.** Box plot showing the high degree of reproducibility of the dbscan clustering presented in Fig 4A. The dbscan algorithm was run on the subsample tSNE maps with the same parameters as for the full sample tSNE map (eps = 4.9, minPts = 3). The Jaccard index was calculated for each cluster using the clustering presented in Fig 4A as reference and all indexes were averaged over all clusters to score each of the 10 selected maps, providing 10 scores for each sub-sample. The blue boxes filled in gray represent the interquartile range, the red horizontal lines correspond to the median values, the whiskers cover the 2.7 standard deviation around the mean under a normality assumption, and the red crosses show the outliers beyond the limit of the whiskers.

### Appendix Tables

**Table S1.**
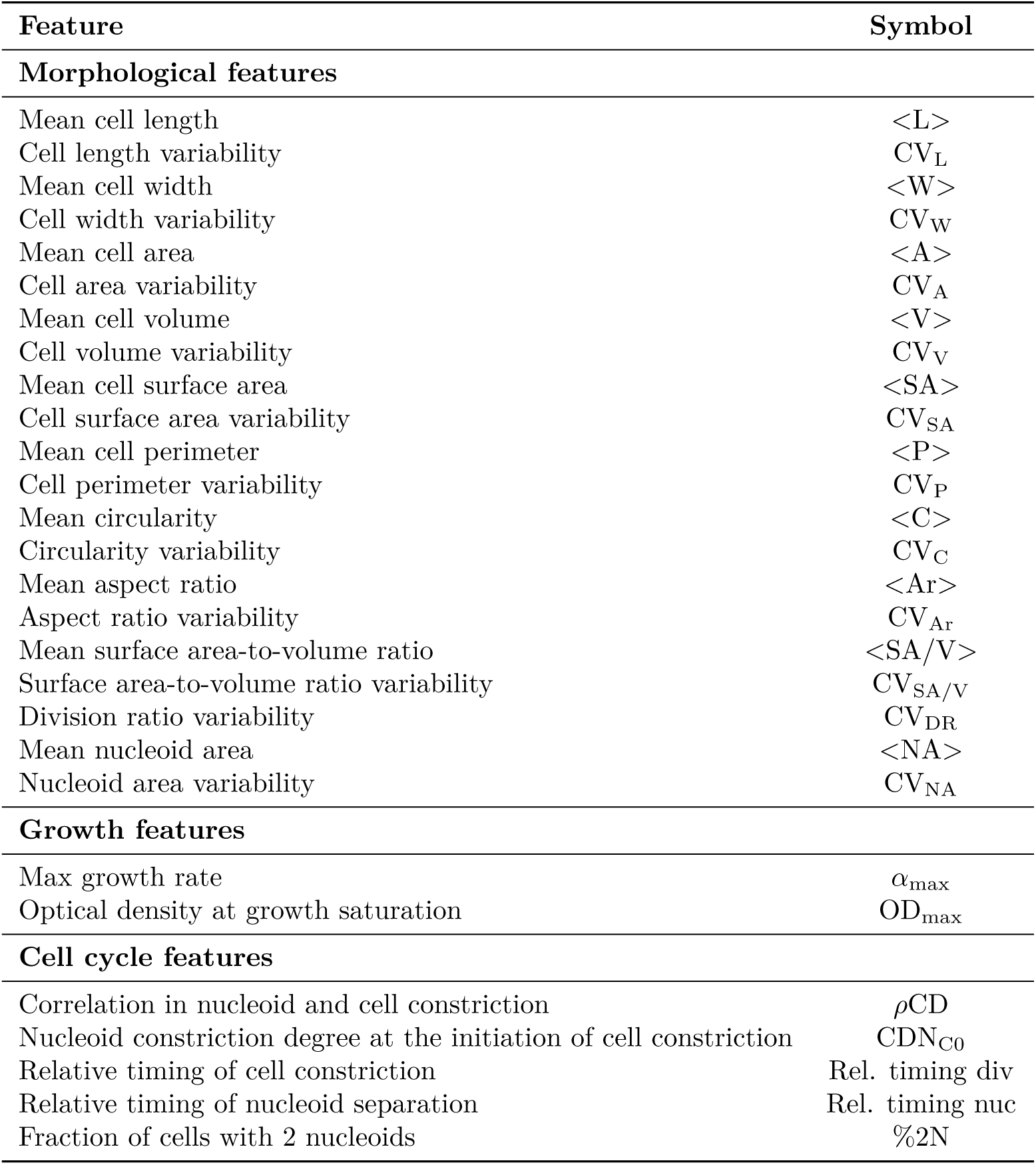
Features considered in this study and their associated symbols. The aspect ratio was defined as the ratio of cell width over cell length at the single-cell level. The circularity, C, was defined as 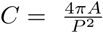, at the single-cell level, where P stands for perimeter and A for area. The relative timing of cell constriction and nucleoid separation were estimated as the proportions of cells without any significant constriction (constriction degree <0.15) or with a single nucleoid, respectively. For all cells with a significant constriction degree, we calculated the Pearson correlation coefficient between the constriction degrees of the cell and of its nucleoid (*ρ*CD). The nucleoid constriction degree at the initiation of cell constriction (CDN_C0_) was determined as the intercept of a line with a slope determined by the correlation coefficient that best fitted the single-cell data used to calculate *ρ* CD (see Appendix Fig S1E).

| Feature | Symbol |
| --- | --- |
| **Morphological features** |  |
| Mean cell length | $\langle L \rangle$ |
| Cell length variability | $CV_L$ |
| Mean cell width | $\langle W \rangle$ |
| Cell width variability | $CV_W$ |
| Mean cell area | $\langle A \rangle$ |
| Cell area variability | $CV_A$ |
| Mean cell volume | $\langle V \rangle$ |
| Cell volume variability | $CV_V$ |
| Mean cell surface area | $\langle SA \rangle$ |
| Cell surface area variability | $CV_{SA}$ |
| Mean cell perimeter | $\langle P \rangle$ |
| Cell perimeter variability | $CV_P$ |
| Mean circularity | $\langle C \rangle$ |
| Circularity variability | $CV_C$ |
| Mean aspect ratio | $\langle Ar \rangle$ |
| Aspect ratio variability | $CV_{Ar}$ |
| Mean surface area-to-volume ratio | $\langle SA/V \rangle$ |
| Surface area-to-volume ratio variability | $CV_{SA/V}$ |
| Division ratio variability | $CV_{DR}$ |
| Mean nucleoid area | $\langle NA \rangle$ |
| Nucleoid area variability | $CV_{NA}$ |
| **Growth features** |  |
| Max growth rate | $\alpha_{\max}$ |
| Optical density at growth saturation | $OD_{\max}$ |
| **Cell cycle features** |  |
| Correlation in nucleoid and cell constriction | ρCD |
| Nucleoid constriction degree at the initiation of cell constriction | $CDN_{C0}$ |
| Relative timing of cell constriction | Rel. timing div |
| Relative timing of nucleoid separation | Rel. timing nuc |
| Fraction of cells with 2 nucleoids | %2N |

**Table S2.** Enriched GO terms in the morpho and cell cycle archipelagos. GO term enrichments were assessed by neighborhoods (see Materials and methods). The table shows the enriched GO terms with a p-value adjusted for a false discovery rate below 0.05. A GO term can be enriched in multiple overlapping neighborhoods. The table lists, for each enriched GO term, the highest (least significant) adjusted p-value and the associated p-value that was calculated for all the overlapping neighborhoods where each GO term was enriched. The corresponding p-value is also reported. The color of the enrichment zone indicated in the “neighborhood” column refers to the color of the neighborhoods where the GO term is enriched in Fig 4E and Fig 5D. Finally, the type of ontology with which each GO term is associated is indicated in the “ontology” column.

| GO term | Highest p-value | Highest adjusted p-value | Neighborhood | Ontology |
| --- | --- | --- | --- | --- |
| Morpho archipelago |  |  |  |  |
| Arginine biosynthetic process | 8.11E-11 | 6.49E-08 | dark blue | biological_process |
| Cellular amino acid biosynthetic process | 3.55E-10 | 2.84E-07 | dark blue | biological_process |
| ATP biosynthetic process | 5.09E-08 | 4.65E-05 | grey | biological_process |
| ATP synthesis coupled proton transport | 2.19E-08 | 2.00E-05 | grey | biological_process |
| ion transport | 4.28E-06 | 3.92E-03 | grey | biological_process |
| plasma membrane ATP synthesis coupled proton transport | 7.32E-09 | 6.69E-06 | grey | biological_process |
| proton transport | 1.93E-06 | 1.76E-03 | grey | biological_process |
| proton-transporting ATP synthase activity, rotational mechanism | 2.19E-08 | 2.00E-05 | grey | molecular_function |
| proton-transporting ATPase activity, rotational mechanism | 2.19E-08 | 2.00E-05 | grey | molecular_function |
| biotin biosynthetic process | 3.89E-05 | 4.14E-02 | light blue | biological_process |
| FtsZ-dependent cytokinesis | 5.96E-06 | 4.24E-03 | orange | biological_process |
| cell division | 1.32E-05 | 2.82E-02 | red | biological_process |
| cysteine biosynthetic process | 3.70E-05 | 4.74E-02 | slate blue | biological_process |
| hydrogen sulfide biosynthetic process | 1.11E-06 | 1.02E-03 | slate blue | biological_process |
| sulfate assimilation | 2.50E-07 | 2.29E-04 | slate blue | biological_process |
| enterobacterial common antigen biosynthetic process | 5.03E-05 | 4.60E-02 | yellow | biological_process |
| Cell cycle archipelago |  |  |  |  |
| hydrogen sulfide biosynthetic process | 2.75E-06 | 2.09E-03 | blue | biological_process |
| sulfate assimilation | 1.16E-05 | 8.76E-03 | blue | biological_process |
| sulfite reductase complex (NADPH) | 4.83E-05 | 3.66E-02 | blue | cellular_component |
| sulfite reductase (NADPH) activity | 4.83E-05 | 3.66E-02 | blue | molecular_function |
| phosphate ion transmembrane transport | 2.75E-06 | 6.26E-03 | brown | biological_process |
| phosphate ion transport | 2.75E-06 | 6.26E-03 | brown | biological_process |
| ATPase-coupled phosphate ion transmembrane transporter activity | 2.75E-06 | 6.26E-03 | brown | molecular_function |
| arginine biosynthetic process | 1.36E-05 | 6.19E-03 | dark green | biological_process |
| cellular amino acid biosynthetic process | 6.11E-05 | 2.32E-02 | dark green | biological_process |
| biotin biosynthetic process | 6.47E-05 | 2.94E-02 | light blue | biological_process |
| ATP biosynthetic process | 4.71E-05 | 2.14E-02 | light green | biological_process |
| ATP synthesis coupled proton transport | 2.70E-05 | 1.23E-02 | light green | biological_process |
| ion transport | 6.47E-05 | 3.68E-02 | light green | biological_process |
| plasma membrane ATP synthesis coupled proton transport | 5.44E-07 | 1.24E-03 | light green | biological_process |
| proton transport | 6.47E-05 | 3.68E-02 | light green | biological_process |
| proton-transporting ATP synthase complex, coupling factor F<sub>o</sub> | 7.70E-06 | 1.75E-02 | light green | cellular_component |
| hydrogen ion transmembrane transporter activity | 7.70E-06 | 1.75E-02 | light green | molecular_function |
| proton-transporting ATP synthase activity, rotational mechanism | 2.70E-05 | 1.23E-02 | light green | molecular_function |
| proton-transporting ATPase activity, rotational mechanism | 2.70E-05 | 1.23E-02 | light green | molecular_function |
| enterobactin biosynthetic process | 7.70E-06 | 3.51E-03 | orange | biological_process |
| 2,3-dihydroxybenzoate-serine ligase activity | 7.70E-06 | 3.51E-03 | orange | molecular_function |

### Appendix Parameters

**Parameters S1. Parameters for cell identification in MicrobeTracker**. The parameters listed below were used to generate cell outlines in MicrobeTracker (Sliusarenko et al, 2011). These parameters can be copied and pasted into the MicrobeTracker parameter panel.

% This file contains MicrobeTracker %settings optimized for wildtype E. %coli

%cells at 0.064 um/pixel resolution %(using algorithm 4)

algorithm = 4

%Parallel Computation

runSerial = 0

maxWorkers = 12

%splitRegions

displayW = 0

wShedNum = 3800

% Pixel-based parameters

getmesh = 1

Nkeep = 320

areaMin =300

areaMax = 3000

scaleFactor = 1

thresFactorM = 0.974

thresFactorF = 0.974

splitregions = 1

edgedetection = 1

edgemode = 1

edgeSigmaL = 1.5

logthresh = 1

edgeSigmaV =0.5

valleythresh1 = 0.0002

valleythresh2 = 1

crossthresh = 0.15

repCoeff1 = 0

attrCoeff1 = 0

erodeNum = 0

opennum = 6

threshminlevel = 0.70

% Constraint parameters

fmeshstep = 1

meshstep = 1

cellwidth = 15

fsmooth = 100

imageforce = 8

wspringconst = 0

rigidityRange = 2.5

rigidity = 1

rigidityRangeB = 8

rigidityB = 5

attrCoeff = 0.2

repCoeff = 0.6

attrRegion = 4

horalign = 0.2

eqaldist = 2.5

% Image force parameters

fitqualitymax = 0.5

forceWeights = [0.25 0.65 0.25]

dmapThres = 2

dmapPower = 2

gradSmoothArea = 0.5

repArea = 0.9

attrPower = 4

neighRep = 5

% Mesh creation parameters

roiBorder = 22.5

noCellBorder = 2

maxmesh = 1000

maxCellNumber = 2000

maxRegNumber = 10000

meshStep = 1

meshTolerance = 0.01

meshWidth = 16

% Fitting parameters

erodeNum = 1

fitDisplay1 = 0

fitDisplay = 0

fitConvLevel = 0.26

fitMaxIter =500

fitMaxIter1 =500

moveall = 0.1

fitStep = 0.2

fitStepM = 0.6

% Joining and splitting

splitThreshold = 0.4

joindist = 5

joinangle = 0.2

joinWhenReuse = 1

split1 = 1

% Other

bgrErodeNum = 4

sgnResize = 1

aligndepth = 1

**Parameters S2. Parameters for nucleoid identification in Oufti**. The parameters listed below were used to generate nucleoid outlines using the objectDetection module in Oufti (Paintdakhi et al, 2016).

|  |  |
| --- | --- |
| Background subtraction method | 3 |
| Background subtraction threshold | 0.1 |
| Background filter size | 8 |
| Smoothing range (pixel) | 3 |
| Magnitude of LOG filter | 0.1 |
| Sigma o PSF | 1.62 |
| Fraction of object in cell | 0.4 |
| Minimum object area | 50 |

